# Combinatorial metabolic engineering of alkane biosynthesis in the osmotolerant yeast *Debaryomyces hansenii* CBS 767

**DOI:** 10.1101/2025.10.23.684212

**Authors:** Zekun Li, Nilesh Kumar Sharma, Sarah Weintraub, Eric Young

## Abstract

*Debaryomyces hansenii* is a promising yeast with diverse potential applications, one of which is producing oleochemicals from inexpensive biomass. Yet, combinatorial metabolic engineering in *D. hansenii* is not possible because, like many nonconventional organisms, key genomic data and genetic elements are missing. Here, we report phenotypic characterization, genomic integration loci, and modular genetic parts that together enable combinatorial metabolic engineering of an alkane pathway in *D. hansenii.* Phenotypic characterization revealed that *D. hansenii* produces lipids when grown on components of lignocellulosic and algal biomass in standard and saline media. Notably, *D. hansenii* produced 57.25% more lipids than *Yarrowia lipolytica* Po1f when grown on glucose. We designed genomic integration loci and derived gene expression elements from a resequenced *D. hansenii* CBS767 genome. While characterizing the integration sites, we also optimized the transformation procedure, validated selection markers, and determined fluorescent reporters. Using homology to known elements in *S. cerevisiae*, we derived 23 promoters and 24 terminators and made them compatible with a modular cloning standard so pathway constructs could be made with automated liquid handling. Flow cytometry measurements show that the promoters span an expression range of three orders of magnitude and the terminators span one order of magnitude. The new part collection was used to construct a genomically integrated combinatorial library of 18 alkane biosynthesis pathways. With no other genetic modifications, the best strain produced 38 mg/L of heptadecane. This is the highest titer observed in a microbe with no other modifications besides the biosynthetic genes. Thus, this work establishes *D. hansenii* as a cell factory for synthesizing oleochemicals from biomass while providing a blueprint for leveraging phenotyping, genomics, modular parts collections, and automated liquid handling for making cell factories from nonconventional organisms. Furthermore, the genetic parts collection now enables functional genetics for various applications in *Debaryomyces* yeasts.

## INTRODUCTION

Yeasts are effective cell factories for production of foods and chemicals and have been in use for centuries. The post-genomic era and the advance of genome editing has opened new frontiers for yeasts as designer cell factories. Specifically, there has been much interest in yeasts that overproduce fatty acids, since they are precursors for diverse oleochemicals with myriad industrial and commercial uses (Blazeck et al., 2013; Runguphan and Keasling, 2014; Xu et al., 2016; Zhou et al., 2016; Zhu et al., 2017). However, the scale and economics of these markets has so far limited commercial adoption because sugar costs are expensive relative to the products. Inexpensive, abundant biomass like lignocellulose and algae has long been proposed as an ideal carbon source, yet depolymerization of this biomass yields significant amounts of pentose sugars and sugar alcohols that are not efficiently fermented by many yeasts (Subtil and Boles, 2011; Ruchala and Sibirny, 2021). This remains the case despite extensive engineering (Gopinarayanan and Nair, 2019; Sasaki and Yoshikuni, 2022).

*Debaryomyces hansenii* has many advantageous characteristics that could make it an ideal platform host for oleochemical production from inexpensive feedstocks. It is naturally halotolerant and xerotolerant, therefore it thrives in saltwater (Aggarwal and Mondal, 2009; Navarrete et al., 2021). It naturally assimilates and ferments D-glucose, D-xylose and L-arabinose, the major monosaccharides in lignocellulosic biomass (Breuer and Harms, 2006). It can also grow on mannitol, a sugar alcohol found in algal biomass (Navarrete et al., 2022). Another remarkable characteristic of *D. hansenii*, shared with other oleaginous yeasts is its capacity to accumulate fatty acids, reaching up to 70% of its dry biomass (Breuer and Harms, 2006). *D. hansenii* is also flavinogenic – it accumulates significant riboflavin, a nutritional vitamin that is produced microbially (Sibirny and Voronovsky, 2009; Dmytruk et al., 2020; Weintraub et al., 2025). Furthermore, *D. hansenii* is Generally Regarded As Safe (GRAS), BioSafety Level 1 (BSL1), haploid, and unicellular. It is classified in the CTG clade of yeasts where the leucine CUG codon is predominantly translated as serine instead of leucine (Papon et al., 2014). This alternative codon usage is shared by many yeast species of biotechnological and biomedical importance such as *Scheffersomyces stipitis*, *Candida albicans*, and *Candida auris* (Cao et al., 2018; Defosse et al., 2018). *D. hansenii* is genetically tractable, and genetic modification with CRISPR has been reported (Spasskaya et al., 2021; Strucko et al., 2021). Therefore, *D. hansenii* could be an ideal host for production of oleochemicals and flavins from high ionic strength mixtures of diverse sugars.

Here, we show that *D. hansenii* has high production of lipids, high production of riboflavin, and fast growth on multiple carbon sources in defined and seawater analog media. This confirms that *D. hansenii* not only grows on pentoses but produces significant amounts of lipids as well. We compare this to two other yeast species: *Saccharomyces cerevisiae* and *Yarrowia lipolytica. S. cerevisiae* is widely used in fermentations, has extensive and sophisticated genetic tools, and well-established genetics; and the oleaginous yeast *Y. lipolytica* is another potential oleochemical platform host with established genetic tools and genomic information (Xu et al., 2016). However, both yeasts cannot consume pentose sugars, and in the case of *S. cerevisiae*, prefer to produce ethanol rather than lipids.

We further develop genome engineering tools in *D. hansenii* by determining integration sites and recoding common antibiotic selection and fluorescent protein reporters to be compatible with the CTG clade codon usage of this yeast. We derive homologs to common *S. cerevisiae* gene expression parts in the *D. hansenii* genome. All of these elements are formatted in a modular genetic part collection. Thus, the collection includes integration sites, recoded selection markers and reporters compatible with CTG clade yeasts, and 47 gene expression elements – 23 promoters and 24 terminators.

We then demonstrate the utility of this modular part collection by performing combinatorial metabolic engineering on a heterologous long-chain alkane pathway for heptadecane production. This pathway consists of a thioesterase (TES), a carboxylic acid reductase (CAR), and an aldehyde deoxygenase (ADO). Several versions of this pathway have been reported. We recoded and tested combinatorially two thioesterases - *Helicobacter pylori* TES (Bi et al., 2016) and *Mus musculus* TES (Chen et al., 2014), three carboxylic acid reductases - *Clavibacter michiganensis* CAR (Gitaitis and Beaver, 1990; Gartemann et al., 2008), *Nocardia lowensis* CAR (He et al., 2004; Venkitasubramanian et al., 2007), and *Mycobacterium marinum* CAR (Zhou et al., 2016), and three aldehyde deoxygenases - *Atelocyanobacterium thalassa* ADO (Lea-Smith et al., 2015), *Nostoc punctiforme* ADO (Zhou et al., 2016), and *Prochlorococcus marinus* ADO (Xu et al., 2016). This totaled 18 different pathways. The best pathway produces 38 mg/L heptadecane, the highest titer of any strain with just these three genes added. Therefore, this work establishes *D. hansenii* as a potential oleochemical cell factory and expands functional genetics in this organism.

## MATERIALS AND METHODS

### Yeast strains and media

*Saccharomyces cerevisiae* S288C (ATCC 204508), *Yarrowia lipolytica* Po1f (ATCC MYA-2613), and *Debaryomyces hansenii* CBS767 (ATCC 36239) were obtained from the American Type Culture Collection (ATCC, Manassas, Virginia). Routine culture of all yeast species was performed in YPD, consisting of 30 g/L yeast extract peptone (Sunrise Science Products #1877-1KG) and 20 g/L D-glucose (YPD, Sigma-Aldrich #G7021). Fermentation media varied carbon source and salinity. The base media were YP, which was 30 g/L yeast extract peptone, or YPS, which was 30 g/L yeast extract peptone and 33 g/L sodium chloride (Sigma-Aldrich, #S3014). The carbon sources included with YP or YPS were either 20 g/L D-glucose called YPD or YPSD, 20 g/L D-xylose (Sigma-Aldrich #X1500) called YPX or YPSX, 20 g/L L-arabinose (Sigma-Aldrich #A3256) called YPA or YPSA, or 20 g/L mannitol (Sigma-Aldrich #A14030) called YPM or YPSM.

### Phenotyping (growth, riboflavin, and total lipids)

Precultures were performed to ensure healthy growing yeast and precondition each strain to each different fermentation media. For seed cultures, single colonies were inoculated into 5 mL YPD in a 15 mL round-bottom tube and shaken overnight at 225 rpm at 25 C in a New Brunswick Innova 44R shaker. Cultures were then diluted 1:1000 into 25 mL fresh YPD medium in a 125 mL culture flask and shaken at 225 rpm at 30C in the Innova 44R until an OD_600_ of at least 1.00 ± 0.02 was reached, as measured by a Genesys 10s UV-VIS spectrophotometer. After seed culture, the cells were transferred to 50 mL conical tubes (Heathrow Scientific #HEA4427B) and pelleted at 500Xg for 5 min at room temperature in an Eppendorf 5810R centrifuge. The supernatant was discarded, and the cells were resuspended in 50 mL water. The cells were then pelleted again, resuspended in water, then pelleted a final time. Finally, the cells were resuspended in 5 mL of one of the fermentation media (YPD, YPX, YPA, YPM, YPSD, YPSX, YPSA, or YPSM) in a 15 mL round bottom tube and shaken at 225 rpm at 30C in the Innova 44R overnight.

These cultures were used to inoculate a 96-well black wall plate (Thermo Scientific #165305) for growth curves and phenotypic analysis on a BioTek Synergy H1M plate reader. First, 90 µL of fermentation media (either YPD, YPX, YPA, YPM, YPSD, YPSX, YPSA, or YPSM) was added to the plate. Then, each culture was diluted with sterile water to an OD_600_ of 1.00 ± 0.02 and 10 µL of each culture was added to the wells containing fermentation media. This yielded an initial OD_600_ of 0.1 in each well of the plate. In the plate reader, cultivation was carried out with reciprocal shaking (120 rpm) at ambient temperature for 48 hours.

For growth, optical density at 600 nm (OD_600_) was measured every 5 minutes. Simultaneously, riboflavin was measured by fluorescence using 440 nm excitation and 520 nm emission. Riboflavin concentration was calculated via the standard curve in Supplemental Figure S1.

At the end of the 48 hour plate reader cultivation, total lipids were measured by Nile red assay. Nile red dye (ThermoFisher #N1142) at a concentration of 0.05 mg/mL was dissolved in DMSO 1:1 v/v. Then, 20 µL of this solution was added to each well of the black wall plate and was given 5 minutes to react. Then, each sample was measured in the BioTek H1M by fluorescence with excitation at 530/25 nm and emission at 590/35 nm and the optical position is set to the top 50% of the well. The kinetic reading was performed for 60 min with 60 sec intervals. The fluorescence measurement was normalized by the OD_600_ to control for variations in cell density between wells. Therefore, total lipids were calculated as the relative fluorescence units per unit of optical density (RFU/ OD_600_).

### Statistical analysis

All quantitative data were calculated as the mean value with corresponding standard deviation obtained from replicates of three independent experiments. The statistical analysis of riboflavin and total lipids were performed using the student’s t-test, and a p-value < 0.05 was considered significant.

### Minimum inhibitory concentration assay (MIC)

A minimum inhibitory concentration (MIC) assay was used to confirm and determine appropriate antibiotic selection markers for this organism. MIC is defined as the lowest concentration of antifungal agents sufficient to inhibit 80% growth compared to control. To do this, a preculture of *D. hansenii* was grown overnight on 5 mL YPD at 25 °C and diluted with YPD to an OD_600_ = 1.0. 100 µL of the preculture was pipetted into each row of columns 2-11 of a 96-well round bottom plate (Corning, 3596). These ten columns were for serial dilution of antibiotics, while the tenth was the no-antibiotic positive control. A negative control of 100 µL YPD was pipetted into column 12. The antifungals hygromycin (ThermoFisher, 10687010), nourseothricin (Jena Bioscience, AB-101L), zeocin (Jena Bioscience, AB-103S), and geneticin (ThermoFisher, 10131-035) were chosen for the experiment as these are common yeast selections, and some have been used in *D. hansenii* (Minhas et al., 2009; Hegemann and Heick, 2011). Stocks of each antifungal were made in concentrations of 5120 µg/mL and 7680 µg/mL. Replicates of antifungal concentration were made by pipetting 200 µL of an antifungal into three rows of column 1 of the 96 well plate (with cells in columns 2-11 and YPD in column 12). A multichannel pipette set to transfer 100 µL was then used to perform a serial dilution from column 1 to column 10. At column 10, 100 µL of the culture and antibiotic mixture was removed to leave 100 µL in every well of the plate. The plate was then incubated at 25 °C for 48 hours on the BioTek Synergy H1M and OD_600_ of every well was measured with column 12 as the blank. Growth measurements were used to calculate the MIC of each antifungal agent.

### Plasmid designs for modular cloning

All plasmids used in this study are summarized in Supplementary Table S3 and the maps are available in the Supplementary Figure S2. Each plasmid used in this work is compatible with the modular cloning standard. The Level 0 part plasmids (pJHC07AB-Pro, pJHC07BC-ORF, pJHC07CD-Ter) and the Level 1 unit plasmids (pJHC15DV-EF, pJHC15DV- FG, pJHC15DV-GH, and pJHC15DV-HI) were previously reported (Collins et al., 2021). Unique Level 2 plasmids with homology to *D. hansenii* intergenic regions flanking the TypeIIS enzyme restriction sites were constructed by Gibson assembly. Of these, two plasmids are for inserting two transcription units – one fluorescent protein and one selection marker (pNKS227_9EG, and pZL227_13EG) – and two plasmids are for inserting four transcription units – the three gene alkane metabolic pathway and one selection marker (pNKS227_9EI, and pZL227_13EI). These are all kanamycin resistant, and each consists of the *ccdb* gene flanked by BbsI sites for “cloning in” a set of transcription units, which in turn is flanked by 5’ and 3’ *D. hansenii* genome homology. This entire sequence is itself flanked by BsaI restriction sites so that a linear fragment consisting of the payload and homology arms may be digested from the Level 2 plasmid.

### Genes and gene expression parts

All sequences of promoters and terminators were derived from the *D. hansenii* genome based on homology to known promoters and terminators from *S. cerevisiae.* Their names are thus derived from that homology (*i.e.* P_act1_ is the promoter sequence derived from the *S. cerevisiae ACT1* homolog in *D. hansenii*). These were derived and modified to eliminate BsaI and BbsI restriction enzyme sites by the Joint Genome Institute, Berkeley, CA, United States as part of their Synthetic Biology Community Science Program (CSP). All promoter sequences are reported in Supplemental Table S4. All terminator sequences are reported in Supplemental Table S5.

Fluorescent reporter gene sequences for mCherry, mCardinal, and GFP were sourced from those commonly used in *Saccharomyces cerevisiae* and recoded for use in *D. hansenii* and with the modular cloning system. Similarly, the SAT1 gene for nourseothricin resistance was sourced from *Candida albicans* and optimized for *D. hansenii* and for compatibility with the modular cloning system. All pathway gene sequences were obtained from various sources (see Supplemental Table S3) and recoded for *D. hansenii.* All recoded reporter, selection, and pathway sequences are reported in Supplemental Table S6.

### Automated modular cloning

Since every part in the collection was cloned into a Level 0 plasmid (Supplemental Figure S3), any arbitrary transcription unit could be assembled. To efficiently assemble gene expression element transcription units and, subsequently, metabolic pathways, we used automated liquid handling to perform the cloning. *E. coli* strains containing Level 0 plasmids were inoculated into the appropriate selection media, grown overnight, and then miniprepped (Qiagen #27104). The concentration of each plasmid was diluted to 20 fmol/μL, and this solution was aliquoted into a Beckman Coulter Echo 384-Well Low Dead Volume source plate (Beckman Coulter #17311983) at 14 μL per well. Then, a script describing the pipetting steps to combine 0.1 μL of each promoter, gene, terminator, and Level 1 destination vector solution into a 96 well PCR plate was programmed into the Echo 525 Plate Reformat software. After dispensing the part combinations, DNA assembly master mix was then added to each well using an Eppendorf EpMotion deck-based pipetting robot. The master mix was scaled based on the following proportions per assembly: 0.2 μL of assembly enzyme, 0.08 μL of T4 ligase, 0.2 μL of ligase buffer, and 0.12 μL of nuclease free water. This totals a 1 μL total reaction volume for each assembly. The PCR plate containing the completed assembly mixtures was then transferred to a thermocycler programmed to run 30 cycles of the following pattern: 37 °C for 2 minutes, 16 °C for 2 minutes, then 50 °C for 5 minutes, ending with 10 minutes at 80 °C to inactivate the enzymes. Assemblies were stored at 4 °C until transformation into *E. coli.* Automated transformation of *E. coli* was performed on the Eppendorf EpMotion as previously described (Keating and Young, 2023). Colonies were picked from the transformation plates and sequence validated by Sanger sequencing (Quintara Bioscience). After Level 1 plasmids were validated, they were subsequently assembled into Level 2 plasmids following the same procedure. These Level 2 plasmids were used for integration into the *D. hansenii* genome.

### Yeast transformation

The method for transformation of *D. hansenii* was modified from methods previously reported (Spasskaya et al., 2021) and is depicted in Supplemental Figure S2. Cells were streaked on YPD-agar plate from glycerol stock and incubated for 2 days at 25 °C. One colony from the plate was picked to start a preculture in 5 mL YPD media and grown at 25 °C overnight. The preculture was then diluted in 50 mL YPD media to OD_600_ 0.0125 and grown until and OD_600_ of 2.2-2.6 was reached. Cells were pelleted at 3200x*g* for 5 min, the media discarded, and the pellet resuspended in 6 mL of 50 mM sodium phosphate buffer with 25mM dithiothreitol (DTT). This solution was put in a 30 °C incubator, shaken for 15 min, then pelleted again. The pellet was then washed with 40 mL water, repelleted, washed again with 1 mL of ice-cold 1M sorbitol, pelleted, and finally resuspended in 200 μL of 1M sorbitol. These competent cells were aliquoted in volumes of 100 μL mixed with 2-4 μg linearized integrative DNA on ice. The solution was then added into cold 2 mm cuvettes and electroporated in Eppendorf eporator #4309000027 (Eppendorf, USA) at 2300V. Immediately after electroporation, the cells were mixed with 1 mL YPD with 100 mM sorbitol and incubated at 25 °C overnight. The recovery culture was then plated on agar plates with selective media and 0.3 M sorbitol, and incubated at 25 °C for 2-4 days.

### Flow cytometry

Expression strengths of promoter and terminator pairs were determined using GFP (recoded mVenus, see Supplemental Table S6) by cloning promoter-GFP-terminator transcription units along with a nourseothricin selection transcription unit into the appropriate integrativ Level 2 plasmid. The method for characterization by flow cytometry was adapted from Kaishima (Kaishima et al., 2016). The yeast transformants were picked into 14 mL Falcon tube containing 2 mL YPD with 10 mg/L nourseothricin. Falcon tubes were incubated overnight at 25 °C with shaking at 225 rpm. Cultures were inoculated with 20 μL of this pre-culture into 5 mL fresh media in triplicate and then grown overnight to late exponential phase. The overnight yeast cultures were diluted 1:10 by adding 20 μL of the culture into a 96-well flat-bottom plate (Millipore sigma #CLS3370) containing 180 μl sterile water.

These 96-well plates were then placed into a Beckman Coulter CytoFlex S VBY flow cytometer and 20,000 events for each sample was collected. Fluorescence intensities calculated using the FlowCal toolkit (Castillo-Hair et al., 2016).

### Alkane production and quantification

The method for heptadecane quantification was adapted from a previously reported method (Khoomrung et al., 2013). Starter cultures were inoculated in with 5 mL YPD overnight. After two days, cells were pelleted and resuspended in 5 mL fresh media with a 500 μL dodecane overlay, sealed with a semipermeable seal, and cultured in a rotary drum at 30 LC for five more days. Tubes were allowed to separate before removing 10 μL of the dodecane layer for analysis. For alkane quantification, cell pellets were collected from 10 mL cell culture and then freeze dried for 48 h. Metabolites were extracted by a 2:1 chloroform:methanol solution. Samples were analyzed by Agilent GC-MS 6890N. The temperature of inlet, mass transfer line and ion source were kept at 250 LC, 300 LC, and 230 LC, respectively. The flow rate of the carrier gas (helium) was set at 1.0 mL/min, and data were acquired at full-scan mode (50–650 m/z). The program was to hold a temperature of 50 LC for 5 min, then ramp to 140 LC at a rate of 10 LC per min, hold for 10 min, then ramp to 310 LC at a rate of 15 LC per min, finally holding for 7 min. Heptadecane titer was calculated from a standard curve made from a commercial alkane mixture (Sigma Aldrich #04070). There were six standards, all in acetone solvent: 0.0 mg/L, 0.125 mg/L, 0.25 mg/L, 0.5 mg/L, 1 mg/L, and 2 mg/L alkanes. The area under the heptadecane peak was measured for each standard, and this was used to calculate the heptadecane concentration of samples from the engineered *D. hansenii* strains.

## RESULTS

### *D. hansenii* can grow on important monosaccharides, making lipids and riboflavin

We first performed a phenotypic analysis of the growth and production properties of *D. hansenii* CBS 767 in synthetic complete media (CSM) and a seawater analog media (SAM) with different carbon sources (Figure 1). The results confirm that *D. hansenii* CBS 767 can grow on pentoses and mannitol. Notably, its growth rate is increased in the seawater analog media. In comparison to wild type strains of *Y. lipolytica* and *S. cerevisiae* (Figure 2a), *D. hansenii* CBS767 grows faster when grown on glucose than *Y. lipolytica* but slower than *S. cerevisiae* (Figure 2b). As expected, *D. hansenii* has a broader carbon source palette than these other strains. In terms of lipid accumulation, *D. hansenii* produces more lipids in all conditions than both *S. cerevisiae* and *Y. lipolytica* (Figure 2c). This is particularly evident in the seawater analog, pointing towards the unique adaptations that *D. hansenii* has for salt tolerance. We also quantified riboflavin production in each of these conditions (Figure 2d). *D. hansenii* once again produced much more riboflavin than the other strains on glucose and mannitol, but it did not accumulate riboflavin when grown on pentoses. Riboflavin is a precursor to FAD cofactor that powers a great deal of redox reactions in metabolism. Yeast pentose metabolism uses a redox intensive pathway feeding into the pentose phosphate pathway. It is possible that riboflavin does not accumulate during growth on pentoses because it is being cycled to cofactors to manage the additional redox burden of the pentose phosphate pathway (Weintraub et al., 2025). Even so, these growth and production experiments confirm that *D. hansenii* is a promising cell factory for production of oleochemicals from inexpensive biomass.

**Figure 1.**
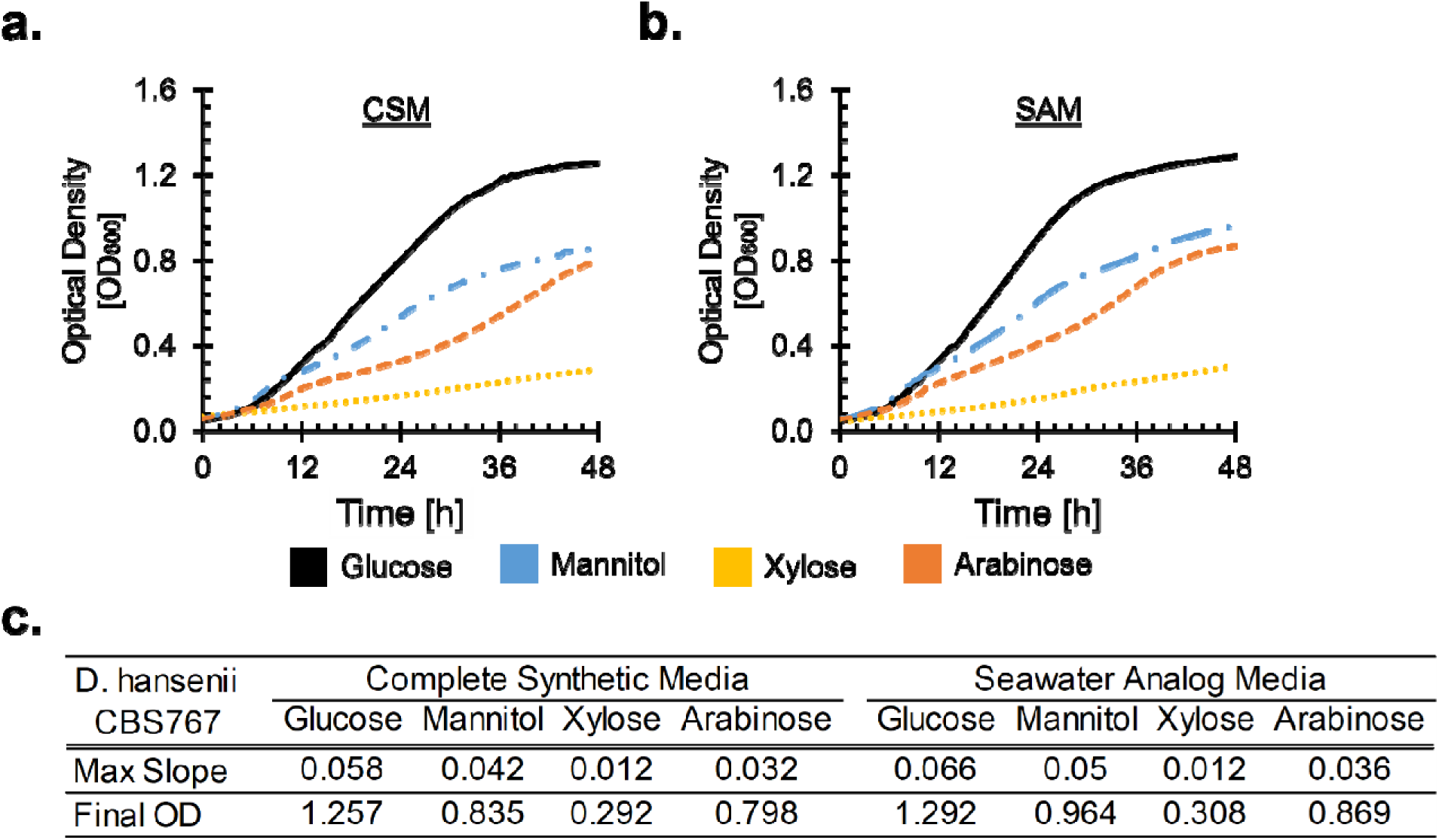
D. hansenii CBS767 and its growth relative to model yeasts. **a.** Growth of *D. hansenii* CBS 767 quantified by optical density at 600 nm (OD_600_) in a plate reader (100 uL media) on complete synthetic media (CSM) with four carbon sources: D-glucose, D-mannitol, D- xylose, or L-arabinose. **b.** Growth of *D. hansenii* quantified by OD_600_ in same plate reader on the same four carbon sources, but in seawater analog media (SAM). **c.** Max slope and final OD_600_ of the growth curves.

**Figure 2.**
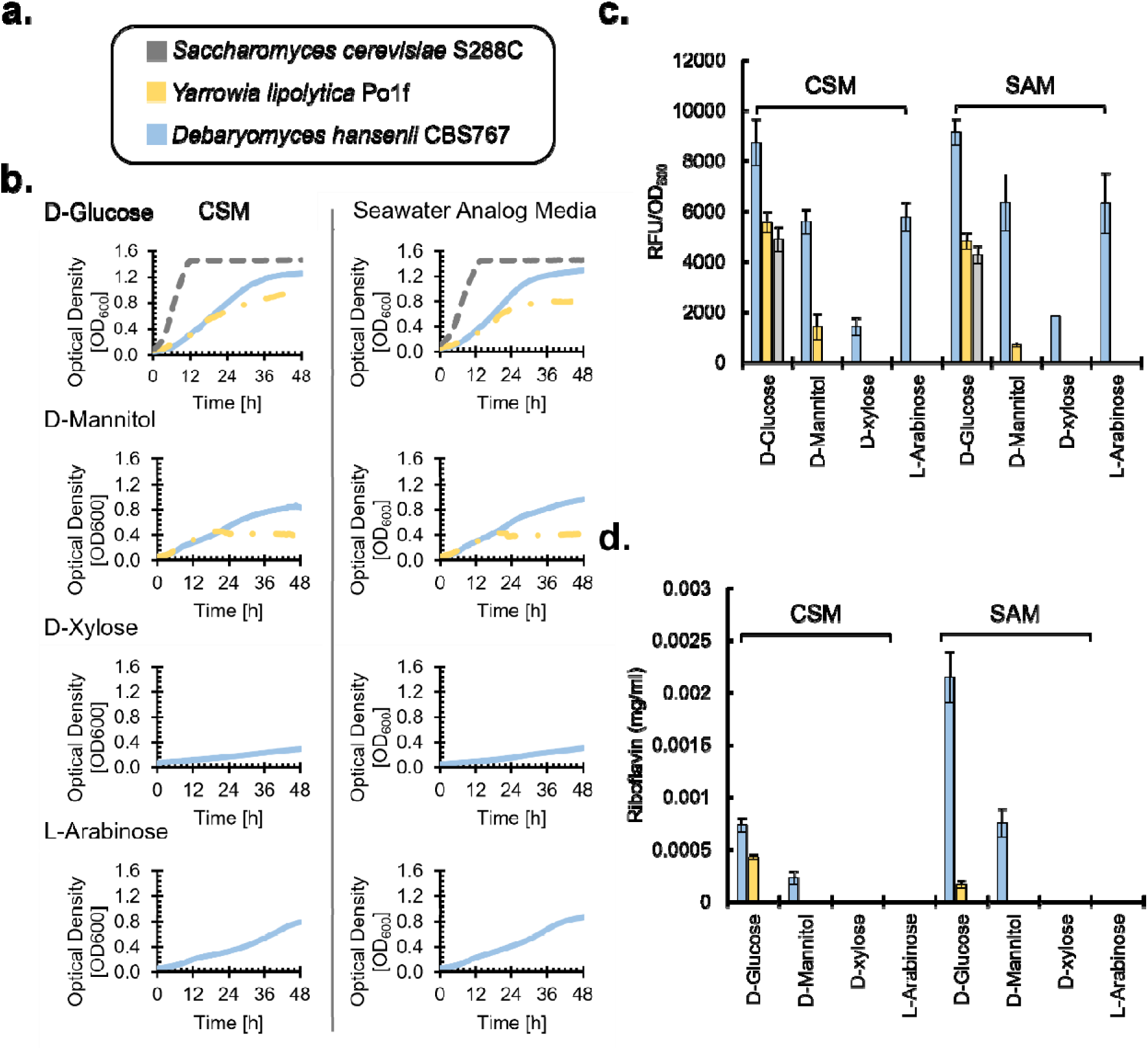
Lipid and riboflavin production of D. hansenii compared to model yeasts. **a.** Color code key for each yeast species in all panels of this figure. **b.** Growth rates of the three yeast species on different carbon sources and CSM or SAM. Max slope and final OD_600_ are reported in Supplemental Table S1 and Supplemental Table S2, respectively. **c.** Lipid productivity of each yeast after two days of growth on the four carbon sources in CSM or SAM quantified by Nile Red staining (relative fluorescence units, RFU, at excitation 530 nm emission 590 nm) divided by cell density (OD_600_). **d.** Riboflavin productivity (mg/mL) of each yeast after two days of growth on four carbon sources in CSM or SAM, quantified by high pressure liquid chromatography (HPLC) correlated to a standard curve (Supplemental Figure S1).

### Genome engineering of *D. hansenii –* Transformation

Successful transformation of *D. hansenii* was difficult in our hands. Therefore, we conducted systematic optimization of an electroporation protocol (Minhas et al., 2009), more fully described in Methods and depicted in Supplemental Figure 2a. In brief, we found that competent cells should be prepared from cells in the mid-log phase, with an optimal OD_600_ reading of around 2.2-2.6. Electroporation with chilled 1M sorbitol and subsequent recovery with YPD with 100mM sorbitol added are crucial steps in the transformation process.

### Genome engineering of *D. hansenii –* Reporters

We then tested fluorescent protein reporters, recoding mVenus green fluorescent protein, and two red fluorescent proteins - mCherry and mCardinal. We designed initial expression cassettes using the known *D. hansenii* promoter P_act1_ (Spasskaya et al., 2021). We integrated these into chromosome E and G of *D. hansenii* (Figure 3a) using a recoded nourseothricin resistance gene as a selection marker and new integrative plasmids for *D. hansenii* (Supplemental Table S3). The results for mCherry, mCardinal, and mVenus are shown in Figure 3 b, c, and d, respectively. We assumed that riboflavin, since it is fluorescent in the GFP range, would significantly interfere with detection of mVenus. However, this was not the case. We found that mCherry and mVenus were both bright reporters, and that the background fluorescence of *D. hansenii* did not greatly interfere with measurement of GFP expression. Therefore, we performed the rest of the experiments with the GFP reporter. Furthermore, in all cases the expression of the fluorescent protein cassettes integrated into chromosome E was stronger than those in chromosome G. Therefore, all subsequent experiments were performed using the integration locus in chromosome E.

**Figure 3.**
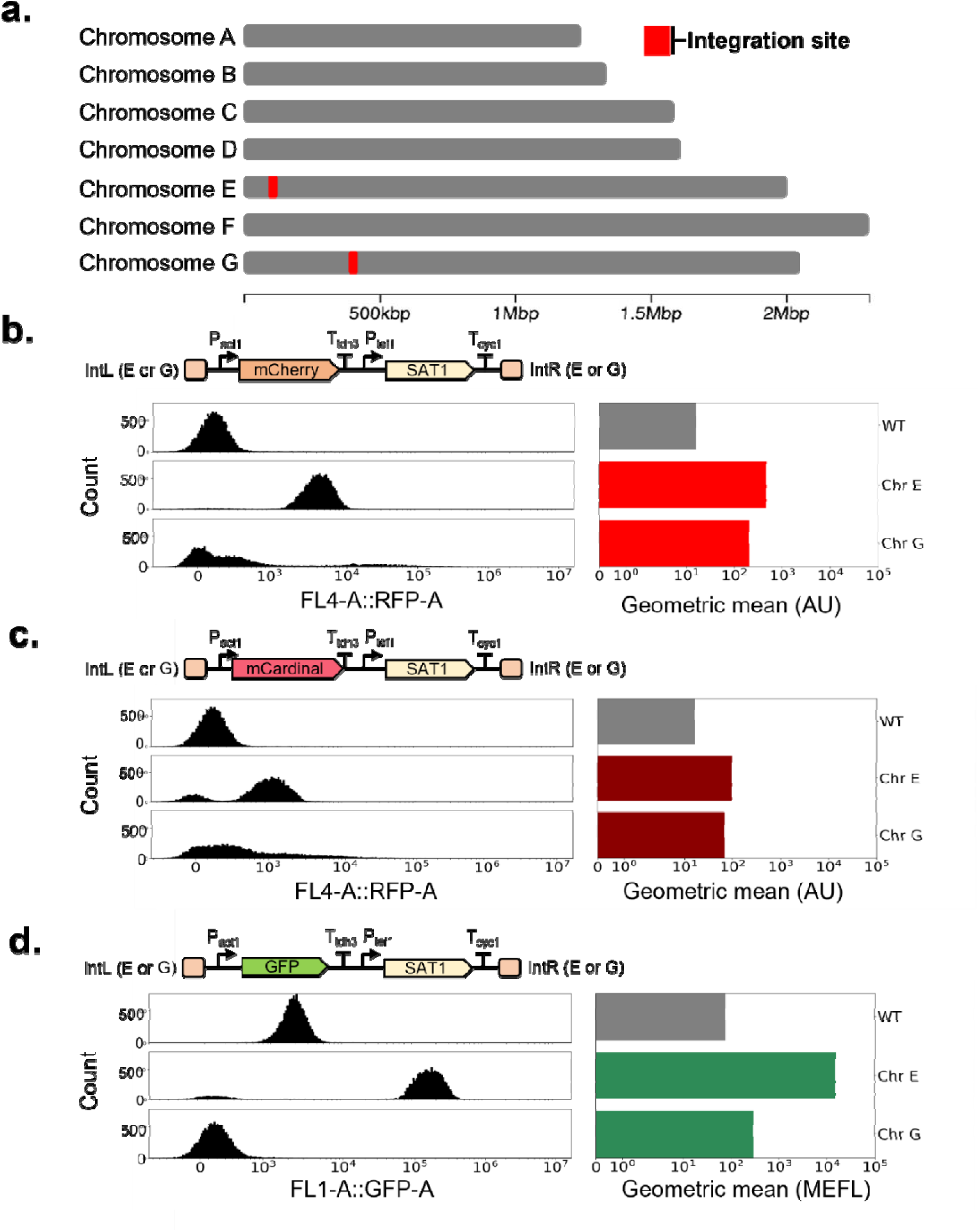
Characterization of integration sites and recoded CTG fluorescent reporters fo *D. hansenii* CBS767. a. Ten possible integration sites across the genome. b. mCherry expression cassette (P_act1_ promoter and T_tdh3_ terminator with nourseothricin resistance, P_tef1_- SAT1-T_cyc1_) integrated into either Chromosome E or G compared to wild type (WT). On the left is a histogram of the FL4 channel area values for 20,000 cells of the population, on the right is a bar chart of the geometric mean fluorescence of the population in arbitrary units (AU). c. Flow cytometry of the cassette with mCardinal compared to WT. d. Flow cytometry of the cassette with GFP compared to WT. On the left is a histogram of the FL1 channel area values for 20,000 cells of the population, on the right is a bar chart of the geometric mean fluorescence of the population in molecules of equivalent fluorescein (MEFL).

### Modular genetic part collection

In order to enable expression of multiple genes at high levels without repeating gene expression elements, we sought to increase the number of promoters and terminators available for *D. hansenii*. Even though *D. hansenii* is an ascomycete yeast related to *S. cerevisiae*, it has been shown that promoters from *D. hansenii* do not function in *S. cerevisiae* (Zeevi et al., 2014). We assumed the reverse was also true, thus instead of directly using known *S. cerevisiae* promoters, we used their sequences to find orthologs in the *D. hansenii* genome with BLAST. The resulting 23 promoters and 24 terminators were synthesized through the DOE Joint Genome Institute Community Science Program to be compatible with our modular cloning system, depicted in Figure 4a. This system uses the same enzymes and scars as a previously published yeast modular cloning (Young et al., 2018; Collins et al., 2021).

**Figure 4.**
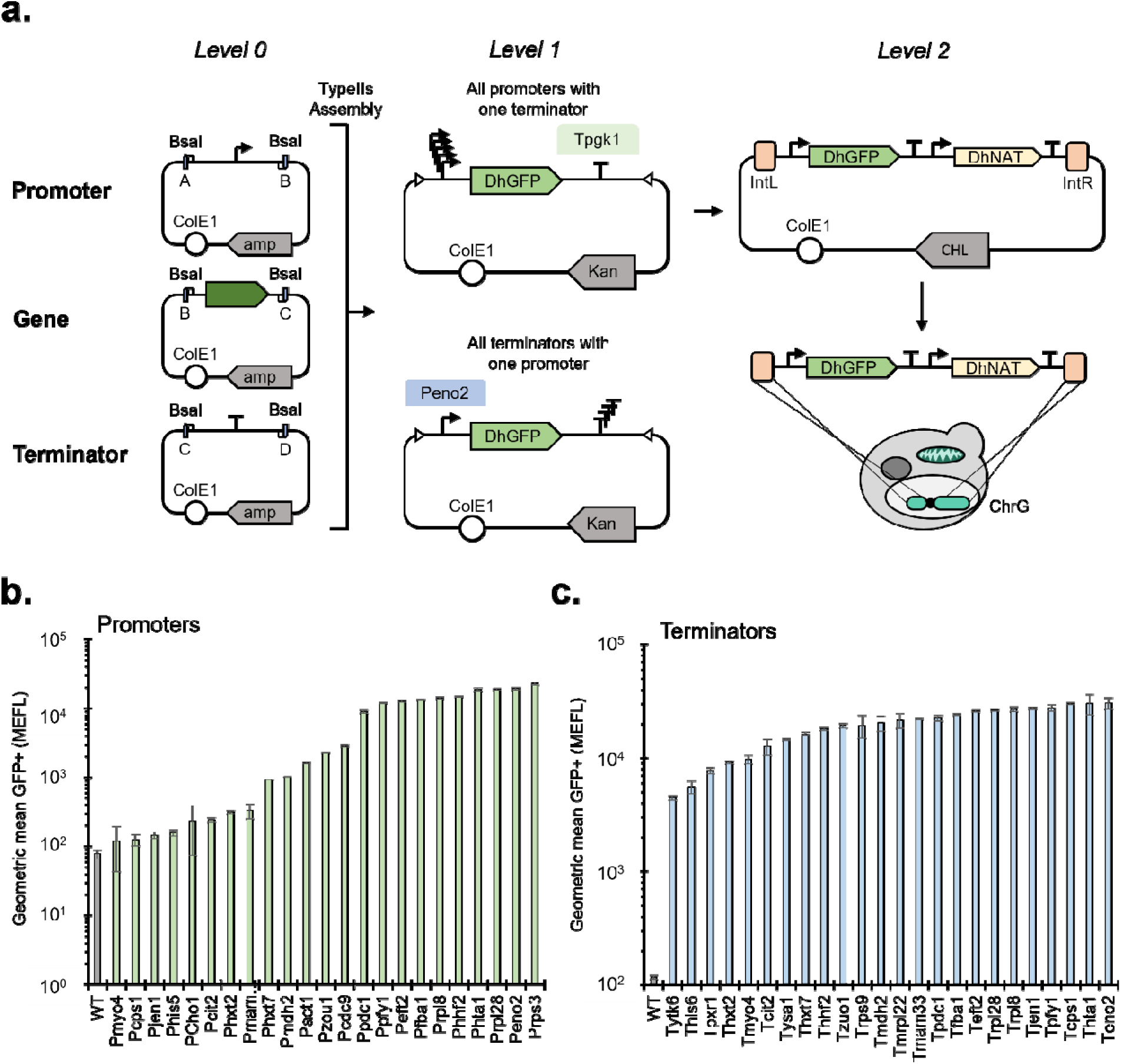
Modular part collection for *D. hansenii* CBS767. **a.** Golden gate assembly and integration scheme for testing gene expression part function. All promoters were assembled with GFP and a standard terminator (T_pgk1_) into a transcription unit that was then combined with a nourseothricin resistance cassette and integrated into chromosome G. All terminators were assembled with GFP and a standard promoter (P_eno1_) into a transcription unit that was then combined with a nourseothricin resistance cassette and integrated into chromosome G. **b.** Geometric mean of flow cytometry data, converted to MEFL, for 23 promoters with T_pgk1_ (green bars) compared to wild type (WT, grey bar). Error bars are the standard deviation in geometric mean for three replicate populations. **c.** Geometric mean of flow cytometry data, converted to MEFL, for 24 terminators with P_eno2_ (blue bars) compared to wild type (WT, grey bar). Error bars are the standard deviation in geometric mean for three replicate population.

We used this modular cloning system in combination with automated liquid handling to construct transcription units for characterizing each promoter and terminator. Each promoter was cloned with two different terminators (T_tdh3_ and T_pgk1_). Each terminator was cloned with two different promoters (P_eno2_ and P_rps3_). This totaled 96 transcription units, which were mixed for assembly using an Echo 525 acoustic liquid handler, transformed via heat shock into *E. coli* using an Eppendorf epMotion liquid handler, and plated onto selective media with the Echo 525 (see Methods).

The expression strength of triplicate transformants was evaluated with flow cytometry using molecules of equivalent fluorescein (MEFL) as a standard unit (Castillo-Hair et al., 2016). The results are shown in Figure 4. When combined with the T_tdh3_ terminator, the promoters P_rpl28_, P_eno2_, P_rps3_, P_eft2_, P_fba1_, P_rpl8_, P_hhf2_, and P_hta1_ showed maximum expression. Medium expression levels were observed for P_mam33_, P_hxt7_, P_mdh2_, P_pfy1_, P_act1_, P_zou1_, P_cdc9_, and P_pdc1_. Promoters P_myo4_, P_cps1_, P_jen1_, P_his5_, P_cho1_, P_cit2_, and P_hxt2_ exhibited low expression levels. Similar expression patterns were observed when combined with the T_pgk1_ terminator, with a few exceptions where different expression levels were observed for certain promoters (Supplementary Information Figure 4). All terminators showed similar expression levels of GFP, but differences in the expression enhancement of terminators are usually only apparent with weaker promoters.

### Combinatorial metabolic engineering for alkane production

With all elements in place, combinatorial metabolic engineering was now possible in *D. hansenii.* To demonstrate this, we designed a three gene pathway for *D. hansenii* that has been shown to produce alkanes in *S. cerevisiae.* Several enzyme variants for this pathway have been published, and they are listed in Supplemental Table S3. We constructed a full factorial set of eighteen pathways consisting of two variants of acyl-CoA thioesterase (TES), three variants of carboxylic acid reductase (CAR), and three variants of aldehyde decarbonylase (ADO) all expressed with unique strong promoters from our collection (Figure 5a). Each of the eighteen variants were assembled by automated liquid handling, transformed, and characterized after 168 hours of growth via GC/MS. A heptadecane peak was obtained at retention time of 23.58 minutes and confirmed by m/z ratio and comparison to a known standard (Figure 5b). All the variants produce heptadecane, but there was a great deal of variance. The results showed that alkane pathway 15, consisting of TES from *Helicobacter pylori*, CAR from *Nocardia iowensis* and ADO from *Prochlorococcus marinus* enabled the highest heptadecane production of 38.31 mg/L (Figure 5c). This represents the highest titer in a yeast species without additional knockouts or overexpressions.

**Figure 5.**
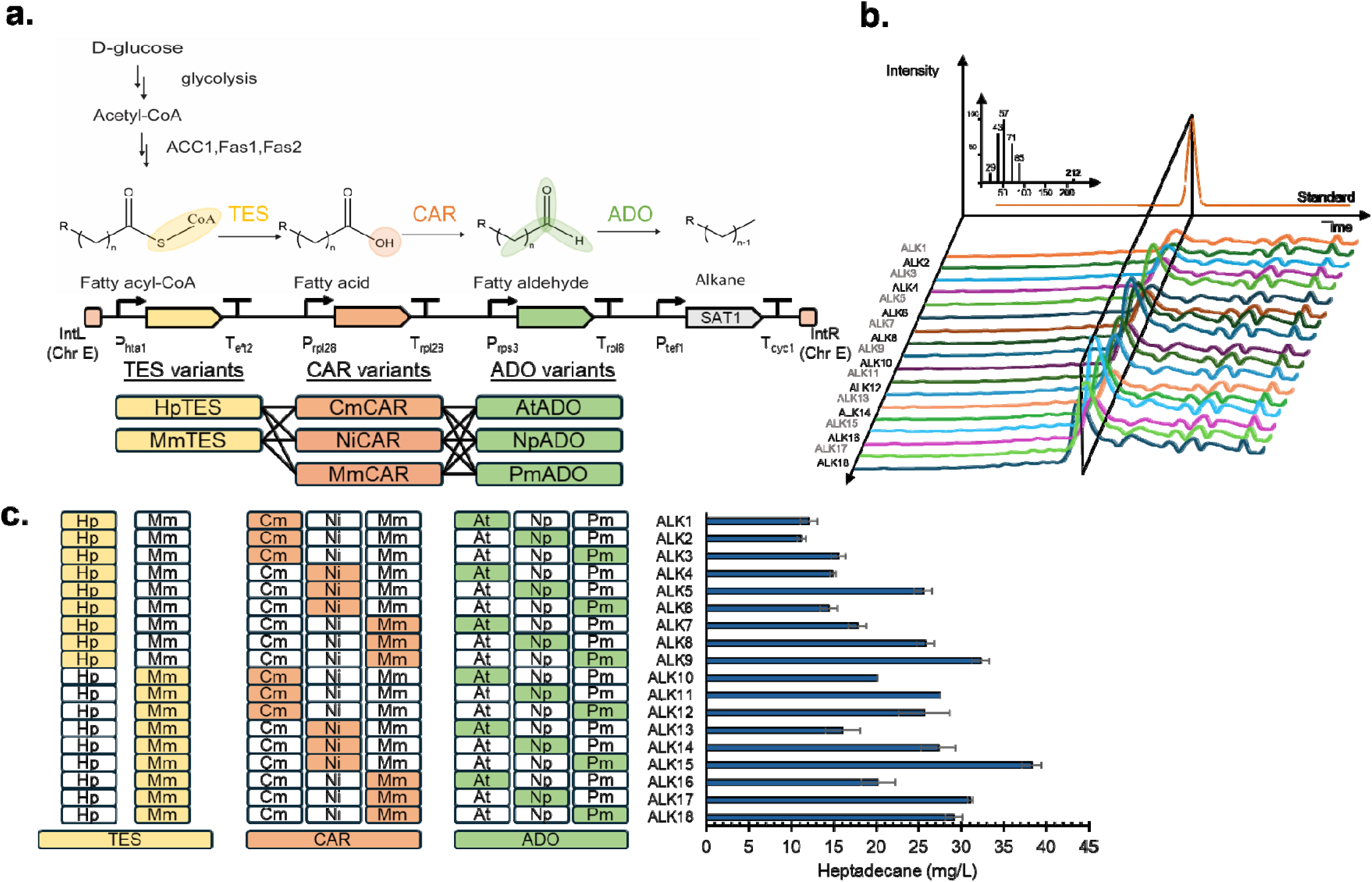
Combinatorial metabolic engineering of alkane biosynthesis in D. hansenii CBS767. a. Simplified metabolic pathway for alkane production. Glucose is converted through glycolysis to acetyl-CoA, which then enters the fatty acid biosynthetic pathway of D. hansenii (acetyl-CoA carboxylase, ACC1, and fatty acid synthases, Fas1 and Fas2). Three heterologous enzymes then catalyze conversion of fatty acids to alkanes. Acyl-CoA thioesterase (TES), carboxylic acid reductase (CAR), and aldehyde decarbonylase (ADO). Pathway design using strong promoters and terminators for combinatorial TES, CAR, and ADO pathway integration. Variants were recoded genes HpTES (Helicobacter pylori), MmTES (Mus musculus), CmCAR (Clavibacter michiganensis), NiCAR (Nocardia iowensis), MmCAR (Mycobacterium marinum), AtADO (Atelocyanobacterium thalassa), NpADO (Nostoc punctiforme), and PmADO (Prochlorococcus marinus). b. Heptadecane identification curve of gas chromatography mass spectrometry (GCMS) quantification..18 alkane extractions in acetone solvent. The area under the curve for all 18 pathways will be converted to mg/L heptadecane using the standard equation. c. Heptadecane titer (mg/L) calculated by a standard curve for the eighteen pathways, and their variant composition. Error bars are the standard deviation of three biological replicates.

## DISCUSSION

In this study, *D. hansenii* was shown to naturally have a desirable combination of fast growth, catabolism of alternative sugars, and elevated biosynthesis of fatty acids. Others have noted that *D. hansenii* has several apparent phenotypic advantages in sugar catabolism, product formation, and tolerance over other nonconventional yeasts (Breuer and Harms, 2006; Prista et al., 2016; Navarrete et al., 2022). Our results corroborate these findings and provide more evidence that *D. hansenii* is of significant interest as a potential cell factory. Like other CTG yeasts, it readily grows on pentoses. Like several oleaginous yeasts it accumulates lipids, but unlike others it does so when grown on pentoses. It grows faster in media with higher ionic strength. It appears to have active cofactor recycling to power redox metabolism that is limiting in other yeasts. This not only supports the case for *D. hansenii* as a promising new host for biomanufacturing, it further argues for more experiments that quantify and compare the phenotypes of potential hosts to determine which organism should be the focus of genetic tool development.

By integrating genomics and genetic engineering, we enabled combinatorial metabolic engineering in this emerging host by developing a modular parts collection. We produced a recoded selection marker, three fluorescent reporters, two integration sites, and 47 gene expression elements. Thus, coupled with other emerging genetic tools like CRISPR (Spasskaya et al., 2021; Strucko et al., 2021), it is now possible to interrogate and manipulate the *D. hansenii* genome on a scale similar to other nonconventional yeasts.

Our results also demonstrate significant production of heptadecane in *D. hansenii* CBS 767. While below the titers of heavily engineered strains, the titer of 38 mg/L we observe for the best pathway is higher than those reported for strains with similar modifications. However, this remains nearly 13% of maximum theoretical yield from glucose on a mass basis. Therefore, this strain represents a starting point for further metabolic engineering.

By demonstrating the favorable phenotypes of *D. hansenii*, developing modular genetic parts, and showing a proof-of-concept for advanced combinatorial metabolic pathway engineering, we show the promise of *D. hansenii* for the production of oleochemicals from renewable resources. Due to the standardized, automated workflow, the combinatorial metabolic engineering strategy demonstrated here may be readily adapted to engineer other valuable products in *D. hansenii* and may also be generalized to other potential yeast cell factories.

## Author Contributions

ZL, NKS, and EMY contributed to the conceptualization of the study. ZL and NKS conducted experiments. SJW performed integration site search and figure preparation. ZL and EMY contributed to manuscript preparation.

## FUNDING

ZL and EMY are supported by National Science Foundation CAREER Award (1944046). NKS was supported by DARPA BioReporters HR0011-19-C-0107. SJW was supported by a National Science Foundation National Research Training (NRT) grant, CEDAR (NRT-HDR-2021871).

## Notes

### Competing Interest Statement

The authors have declared no competing interest.

