## supplemental material for "Combinatorial metabolic engineering of alkane biosynthesis in the osmotolerant yeast *Debaryomyces hansenii* CBS 767"

**Supp. Table S4:** *List of promoter sequences in the parts collection………………………..*	8

**Supp. Table S5:** *List of terminator sequences in the parts collection……………………….*11

**Supp. Table S6:** *DNA sequences of codon-optimized genes used in this study…………..*13

Intensity

| Concentration(mg/ml) | Fluorescence |
| --- | --- |
| 0.01 | 240756 |
| 0.005 | 146528 |
| 0.0025 | 81386 |
| 0.001 | 29649 |
| 0.00025 | 12260 |
| 0.0001 | 3729 |
| 1e-05 | 1972 |
| 0 | 864 |

### Supplemental Figure S1. *Riboflavin standard curve.* There were seven standards for quantification with high pressure liquid chromatography (HPLC), all in acetone solvent: 0 mg/L, 0.0003125mg/mL, 0.000625 mg/mL, 0.00125 mg/mL, 0.0025 mg/mL, 0.005 mg/mL, and 0.01 mg/mL riboflavin.

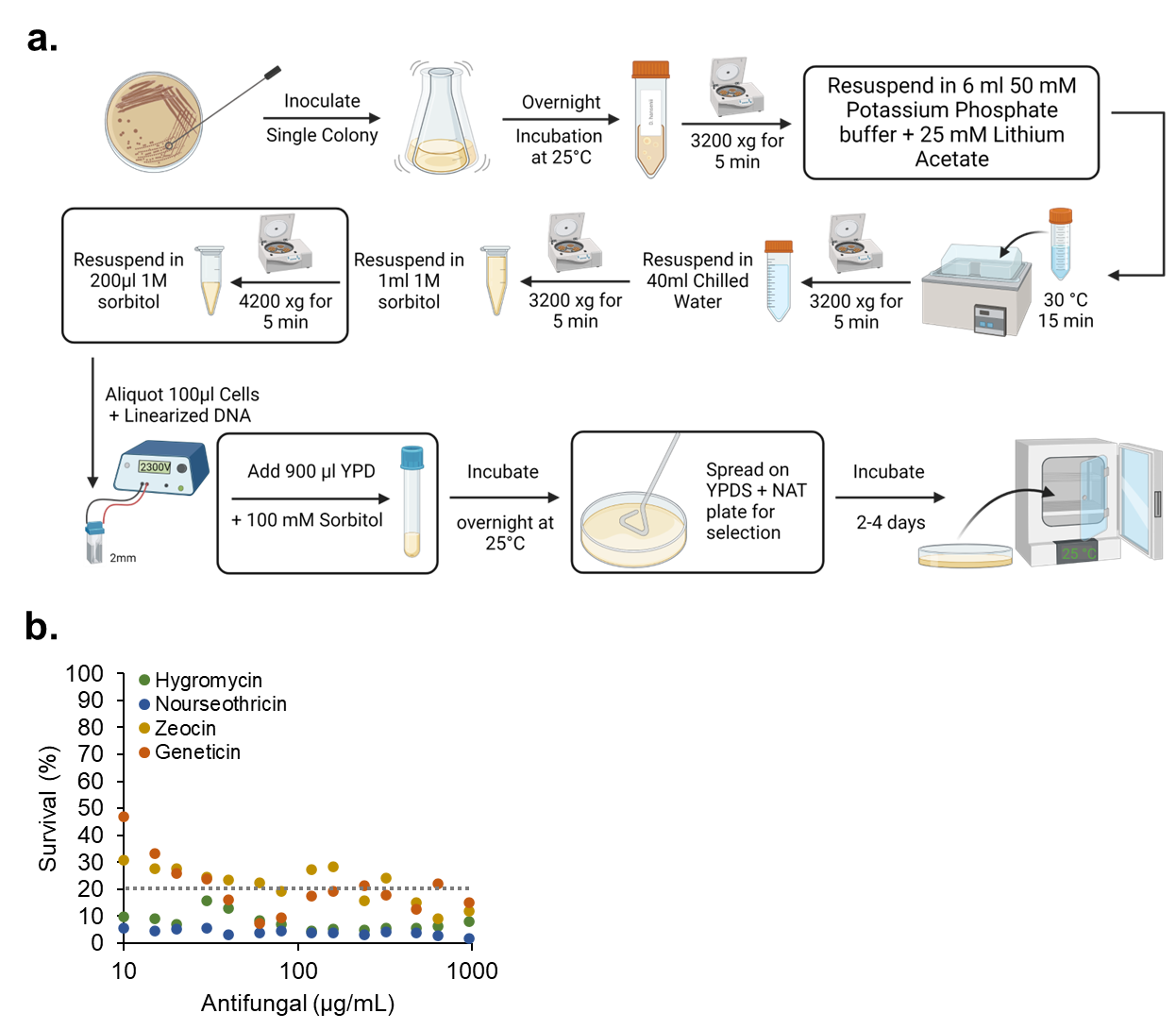

### Supplemental Figure S2. Debaryomyces hansenii *transformation* *optimization.* a. BioRender diagram of the optimized transformation protocol with the optimized steps framed in black outlined boxes. b. Evaluation of the survival percentage of *D. hansenii* on YPD with increasing concentrations of hygromycin, nourseothricin, zeocin, or geneticin. 20% survival, the maximum survival percentage for an antibiotic to be considered effective, is marked as a dotted line. Survival percentages below this line are appropriate for positive selection.

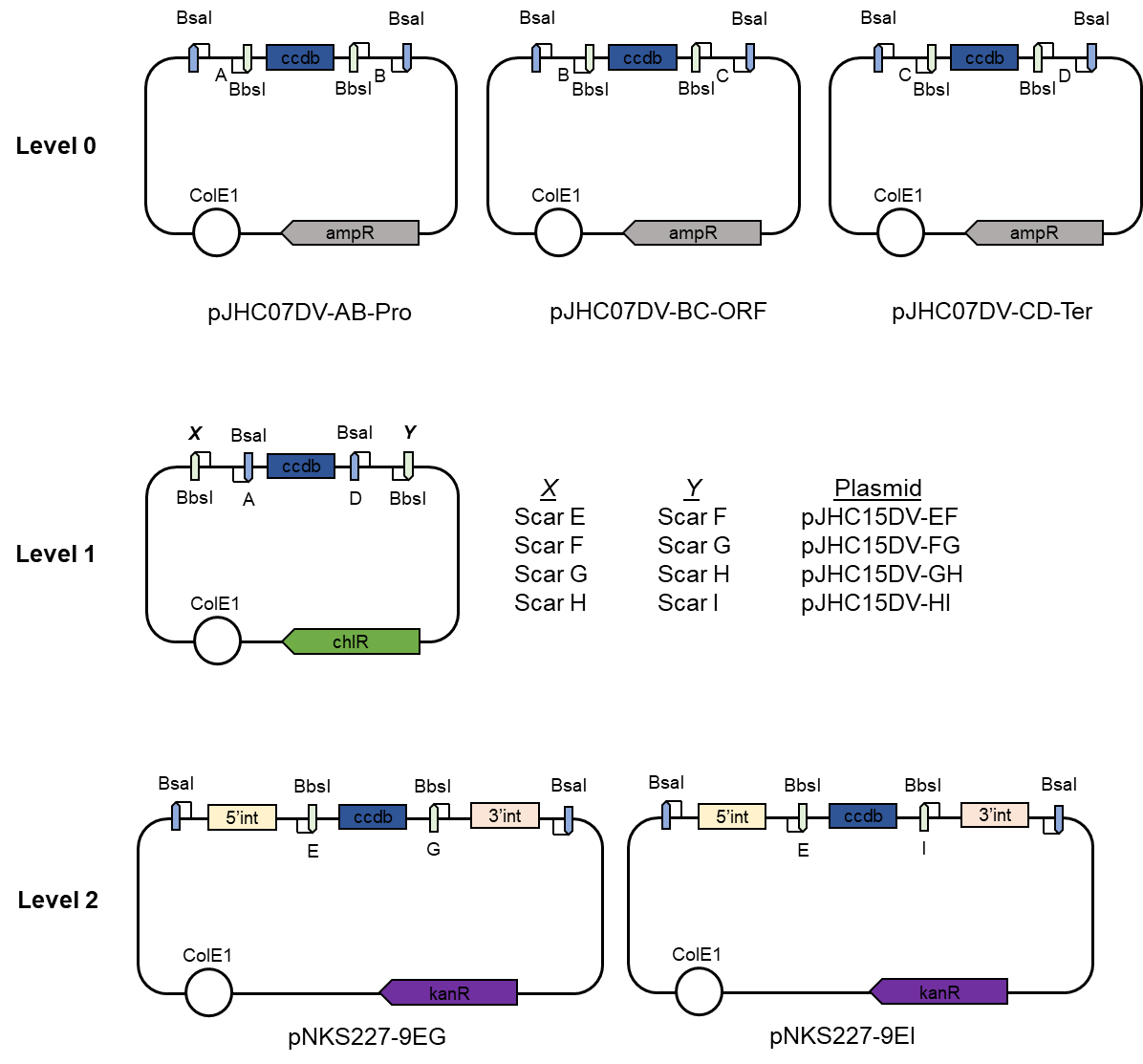

### Supplemental Figure S3: *Yeast modular cloning system for* D. hansenii*.* The hierarchical cloning system uses three Level 0 plasmids for parts, with different scars for promoters (A and B, pJHC07DV-AB-Pro), genes (B and C, pJHC07DV-BC-ORF), and terminators (C and D, pJHC07DV-CD-Ter). Once created, part plasmids can then be combined into a Level 1 transcription unit which has a sequence of scars that dictates assembly order in a pathway (E and F for position 1, F and G for position 2, G and H for position 3, and H and I for position 4). Finally, transcription units may be combined into a full pathway using a Level 2 plasmid that has homology arms to the *D. hansenii* genome (pNKS227-EG for a single gene and selection marker, pNKS227-EI for a 3-gene pathway and selection marker). Once assembled, this fragment may be digested and transformed into the yeast host.

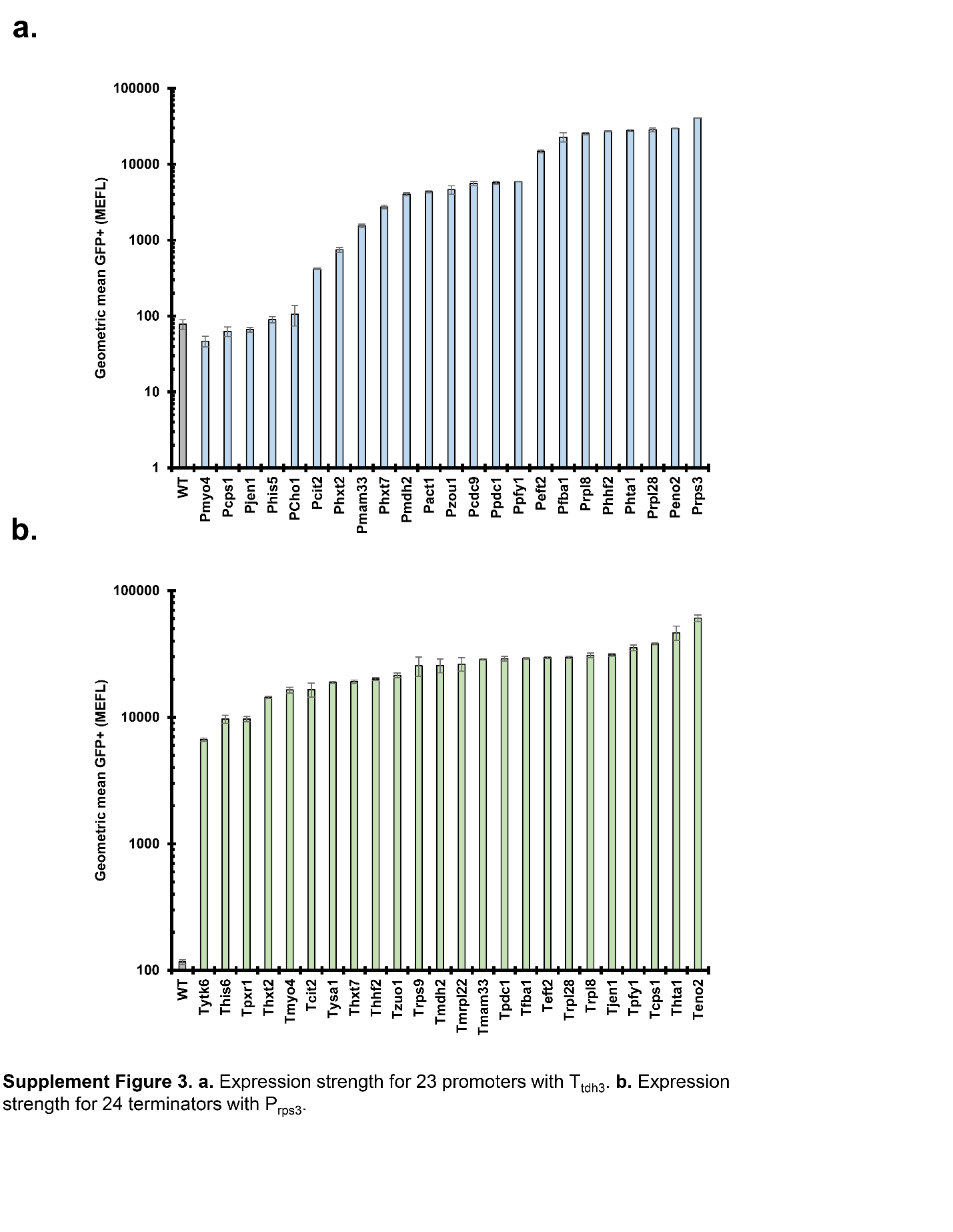

### Supplemental Figure S4. *Gene expression strengths for* D. hansenii *CBS767 parts*. a. Geometric mean of flow cytometry data, converted to MEFL, for 23 promoters with T_tdh3_ (blue bars) compared to wild type (WT, grey bar). Error bars are the standard deviation in geometric mean for three replicate populations. b. Geometric mean of flow cytometry data, converted to MEFL, for 24 terminators with P_rps3_ (green bars) compared to wild type (WT, grey bar). Error bars are the standard deviation in geometric mean for three replicate populations.

### Supplemental Table S1. *Max slope of yeast growth curves*. Growth of *D. hansenii* CBS767*, S. cerevisiae* S288C*, and Y. lipolytica* Po1f quantified by optical density at 600 nm (OD_600_) in a plate reader (100 uL media) on complete synthetic media (CSM) and seawater analog media (SAM) with four carbon sources: D-glucose, D-mannitol, D-xylose, or L-arabinose.

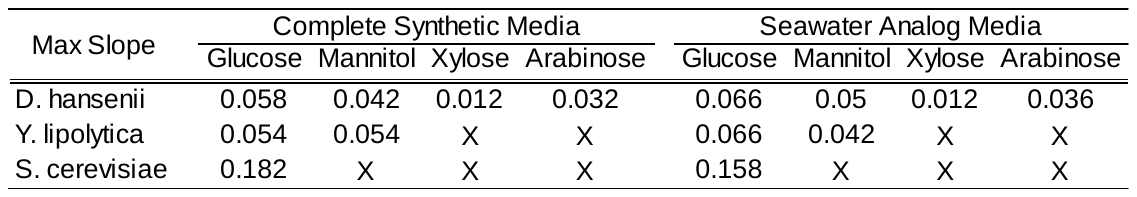

### Supplemental Table S2. *Final OD_600_ of yeast growth curves*. Growth of *D. hansenii* CBS767*, S. cerevisiae* S288C*, and Y. lipolytica* Po1f quantified by optical density at 600 nm (OD_600_) in a plate reader (100 uL media) on complete synthetic media (CSM) and seawater analog media (SAM) with four carbon sources: D-glucose, D-mannitol, D-xylose, or L-arabinose.

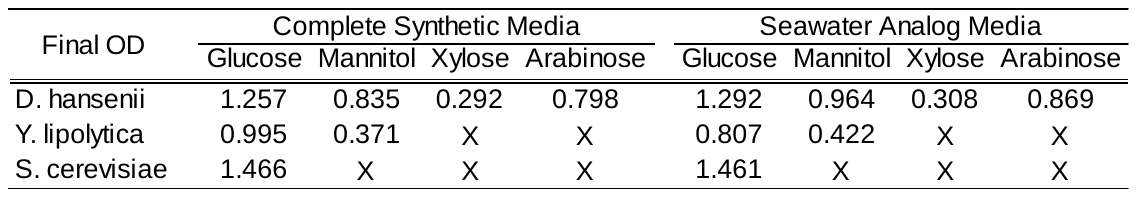

### Supplemental Table S3: *Plasmids used in this study.*

| **Plasmid** | **Description** | **Reference** |
| --- | --- | --- |
| pJHC07AB_PRO | Promoter destination plasmid for modular cloning | (Collins et al., 2021) |
| pJHC07BC_ORF | Open reading frame destination plasmid for modular cloning | (Collins et al., 2021) |
| pJHC07CD_TER | Terminator destination plasmid for modular cloning | (Collins et al., 2021) |
| pJHC15_EF | First position transcription unit destination plasmid | (Collins et al., 2021) |
| pJHC15_FG | Second position transcription unit destination plasmid | (Collins et al., 2021) |
| pJHC15_GH | Third position transcription unit destination plasmid | (Collins et al., 2021) |
| pJHC15_HI | Fourth position transcription unit destination plasmid | (Collins et al., 2021) |
| pJGI07BC_HpTES | *Helicobacter pylori* acyl-CoA thioesterase | This study, gene recoded from (Bi et al., 2016) |
| pJGI07BC_MmTES | *Mus musculus* acyl-CoA thioesterase | This study, gene recoded from (Chen et al., 2014) |
| pJGI07BC_CmCAR | *Clavibacter michiganensis* carboxylic acid reductase | This study, gene recoded from (Gartemann et al., 2008) |
| pJGI07BC_NiCAR | *Nocardia iowensis* carboxylic acid reductase | This study, gene recoded from (He et al., 2004) |
| pJGI07BC_MmCAR | *Mycobacterium marinum* carboxylic acid reductase | This study, gene recoded from (Zhou et al., 2016) |
| pJGI07BC_AtADO | *Atelocyanobacterium thalassa* aldehyde decarbonylase | This study, gene recoded from (Lea-Smith et al., 2015) |
| pJGI07BC_NpADO | *Nostoc punctiforme* aldehyde decarbonylase | This study, gene recoded from (Zhou et al., 2016) |
| pJGI07BC_PmADO | *Prochlorococcus marinus* aldehyde decarbonylase | This study, gene recoded from (Xu et al., 2016) |
| pNKS227_9EG | 2-gene destination vector with Chromosome E homology arms | This study |
| pNKS227_9EI | 4-gene pathway destination vector with Chromosome E homology arms | This study |
| pZL227_13EG | 2-gene destination vector with Chromosome G homology arms | This study |
| pZL227_13EI | 4-gene pathway destination vector with Chromosome G homology arms | This study |

### Supplemental Table S4: *List of promoter sequences in the parts collection.*

| **S/N** | **Promoters** | **Nucleotide sequence** |
| --- | --- | --- |
| 1 | Pdhdhfy1 | TCAGAGCAGATTGTACTGGGTCTCAGTGCTACATGCACGACTCAACTTTCCATACCGGCTGGTTAACCAGATATTAAGATAAAAGCTCTTAGACGCAGCTTTCACTATCATTGCAACGATGATCACATAATTTTTGTAATACAACCCACAGAATCAGAATCACTTCTTGTAAAGCCGTGCTCGGTTGTGGTTTTATGCTACCGCAATCGAATTGCAAAATTTGTACATTAACAAAAATTCTCCAAGAGAAGGATATTATATTTTCACCGAAACTCTAAAATCTTTCTGTTTAACCTCCATATACACATACGTATACATATCGTGTTGTTAAAATACAGTATCGCCTAACGAAGCAAGTTATCGAATAAACCACAAATATGTATACTTCATGTGACCATAGTCACGCGTTTTTGATCGGTCACTATATATGGTCAACAAAATACCAGAAGGATGAAACAACGTTCTTTCTGAATAATAGTAGTAATTTACAGAATTTCACTTAAACAAATACAGTTGAACCATAATAAAGAATGAGAGACCGAGCGGTATCAGCTCACT |
| 2 | Pdhcdc19 | TCAGAGCAGATTGTACTGGGTCTCAGTGCGCGGGTTGGGTGTAGATTAGATTGCGACTTGCTCGCGAACGCTCCGCTACTCCGATTCCGAATGCATCGGAGTTTCTTCGACGGCCCTTGACAGATTTCACTGATATCGTATTCTTACAGCACTTCCTTTCTTCGTTTCCAGAATGTACACACGTATGGCTGTCTTTCCCACCGCAAACATGCGGTTGTTATCTACGGATATGTGGCACCCATGTATCAGATACACCTAAAAGCGCAACGTAGTATTCCGTCGTAATTCTGATTTCGCACTTTTCTGGGAGCCGATGACATGGTCAATTTTTCAGCACACATGGGACATCAGAGGCCATTCTATGGGAAAATACAGCGTTGAGAAGGTGTAAATGGTGTATAATGGACATGCATGTCCTTTAGGAGCCAATTGGTGTATTGTAGGCTTGTTATCGAAAAACACGTAAAAGATTAGTCGCAGGGCCGGAAATTTTTTAGTACGGCCCACTATCAACCGGAGAAATTTTTCATATTTATAGACATAAAAATTTTATATATATAAGGAGTCAAGTCTTTTTAAAATTCAAAGGAAAGCATATTTGTTTATTAAAATATTATACTACATTAGAAAATGAGAGACCGAGCGGTATCAGCTCACTC |
| 3 | Pdheft2 | TCAGAGCAGATTGTACTGGGTCTCAGTGCTTAAGTCCATACACCAACATTCCTACACCAAATTGAATATAAATATGTGCAGGAGCATTCCGCTCTCCTCAAAACCCTGGCACCAAAATGGCTGACATTTCTCCTTCGTTCAACACTATACGGTTCGTTGTTGCTTATTTCCAAGCAAAACTACTCCCAGTAACATGGCCCTCTCTTCTAAATAATATGTAGTCTTAATTGTTCCTATCAGACCTCAGTATCTATCCCAATTTCTCCCATAACCTCATTCCCCGTACTATTGGTTCAATCAAACCCGGGTAACCCAGTCCCAGTATGGTGTTTACTGAGAGAAATATGGGTTACCCGGCGGCTGAAAAATTTTCGATCTCACTTGAGCCATCACCAAAAAATTTTGATACCAATCCCCTCGTGCTGGACAGGTTCCAGTAGCCATTCCCTTGGTTTTTTTTTCCTAGCATTTTATTAAAAAGGATTTTTTAAGATTAGCTAACGTCCAACAACAATAATAATCTATCAAAAATGAGAGACCGAGCGGTATCAGCTCACTC |
| 4 | Pdhfba1 | TCAGAGCAGATTGTACTGGGTCTCAGTGCACCCTCGGACTTCGCCACTAGCCACCTGTATTGCGTCGGAAATCCTACTTGCTGTCCGAAAGTCGTGTTCCCATGAGCGCTCTCTCATTCACTCACTCACATCTCCATAAGCGTCCCGGTGCAATACGGAATAAGCTGGGTGAGTGTAAATAAATTCCGGATGTAAGCTACCATCATTGCTTCAATTGCTACTAAAACCTCGTCATTATAACCAGTCTTATTCAGACGTAACATCATCCGTTTATATAATCGAAGTACGTACATGTAGAGCTATTACGTAGTATAAGGCACGATGACATAATCCATTTCAATAATAAAAGATCATATGGTTCACAGTCTCCCGGGTTAGTAGTGTAGAACTTCGGACTTTTTTCATATTACGTCGAGCACCCCCGAGGGTAGCGTGTTCAAAAACTATATAAGGAGGAGCAAATCACCATTTATTTAATCAATTGGTTTGGTAATATTAATTATACATAAATTATACTAATTAAGAAACAAATGAGAGACCGAGCGGTATCAGCTCACTC |
| 5 | Pdhhhf2 | TCAGAGCAGATTGTACTGGGTCTCAGTGCTTCTTCCCCATAACGTAAACAAACAAACAGTGCGAAAAGCTCATCAGTGCTGTACTGAAAACTGCAACTGCCCCCGGTAATACACAGCACCCCATTTAGACGGTTCGCGACCGCGCAAACGTCACGATCCGCCATTTCTGCTTCCACGCTTAACTTACTCCAGTATTATTGCGACCGGAAAAGACACGATATGGGTGCGAGTAAATGGGTGAGATATTGCCTTTGTTTAGTGTGGAATATGTAATGAGCAAATTGAGATATCTGCTTGAATATTTTTATCATCCCTTATATATATATAAGGCCGCACTTTAAAATTATTAGAAAATTATTCCTTATATTAATATATTTAATTAATATATCTTAATATACTAATATAATAATAATAATATAAAAAATGAGAGACCGAGCGGTATCAGCTCACTC |
| 6 | Pdhrpl28 | TCAGAGCAGATTGTACTGGGTCTCAGTGCCTGCATAGTATTAGACTAAATAAACATCATATATAATTGTAACAAAAATATCTCTATACTACACTCTATGCGTGATTCTGCCGTTAATATGATACAATCTCCAATTAACGAAACAGAATCAAGTAAACTGGCATGATATGACACTATACATGTTTCTATATATCCTTCCATGTACATATCCCGCATTAGGGTTTAGCCCCAATAAACCATACCCAAACAGCATAATGTGCGAAACTCATTCTGTGACTGTTGATGAAAAAAAGCTTTTTTGTGTTATTTTCACTAAGTATGGGAACTTGGTGATGCTCGGTCGCAGTTTGGTGAAAAATTATAGAGTTCAAAAGAATCTTGAAAGTTCTGTTCGTTTACTCTTTATCTAGCTTAATCTATAACATTAATAAATGAGAGACCGAGCGGTATCAGCTCACTC |
| 7 | Pdheno2 | TCAGAGCAGATTGTACTGGGTCTCAGTGCTTACATTGCCAACTACTGGTTTCTCTTCTACTTCTTCCTCGTCTATAACTATTCCAAACTGCCACAATTGGCTCACTTGACTCTATTTCCCTGGTGCTCTACTCCCACTTCCCTATGATTTCCTAATTGCAAAACCACCTCAACAGTCTCACATCATCGTGGTCATAGTGTAGCCATCGCAACCGCCGAGTGTTACGTCATCCCGGAACTCGCAAAGGATGCCCGGGACACTACTGGGCCGTAGAACAGAACTCCGAGCCCGACGTGTCTCGATTGTTGCTCGAATGTACGATAAGTAGTGGAGCTTATTAACCATAACTATACCACCAATTGACGAGCGGTGGCTGACTTAGGCGTGAGAGCATATGGGCTGATCTGTAGACCCGTATATAGACAATAATAAGTGATAGAGCACGTAGAATATGATAGAAAAGACGTAGGATGCGACTCTTTCGGCAGATCGGGAGTGAGTGGTCCACCAACAATAAGTGAAAAATTAATATAAATAGCCAAACTGCACCAAAATTATTCACAATTGACCTGGTGATAATTGATTTATTTTTCATTTGAATACTAAATAAATTAAATATATATATACAAAATGAGAGACCGAGCGGTATCAGCTCACTC |
| 8 | Pdhhxt7 | TCAGAGCAGATTGTACTGGGTCTCAGTGCCGGCCATCGGGTGACAGAAAATTTTATGCACCGGAAAATTTCAAATAATCCGCGATCATCACTAAAATGACACCCGCAGGAAGAATTTTCCGACAACAAGCCGTTGGCTTGAATTTATTTGTTCATCAGCCCATTCTTGATTCTCATTTCACCAGTTTATACAAGAGCAACATTAGGTCGAATATGGAGTGCACGGTGATTAATTTATTAATTTCTTGTATACCTATCTAGATCCATGCTTTAGTATATAAAATGATCTCGGAACATTAAGTAGATGCAGCCGATTAAAAATTATGTGTCCGCTATTTTTCTCCAATAAATCGTTGCCTCATAACTAAGCTGTCCTCAGATTTAGCTATGGTTTAGAACAATGTTCTATGTTGCATGGAGAAACTGGATGTTGATTCGCCTTTTCGCGAACTGAATAATCAGCCTTAGAATGGTATTGTCCCGCACCGGCACTTTTATGAGCGAGTGGGGCAACATGAATTTTTCCGGGGCATGGCACGGAGAAACTGGAGCGTGTATGGGATTAATCTTGCATGCCGGGATAAAAAATTTTTCCAGGTATTTTTCCGGAGGTTCTAAAATCTGTTATGCCTTGATGGTAGAAAGTATAAATAAGTGGGTTATCCCTCTTGAAAATTCAGAATTCTTTTCTTGTATTCAATTATTTATTTAATTTAGAAGTCAAATTATCAATGAGAGACCGAGCGGTATCAGCTCACTC |
| 9 | Pdhhis5 | TCAGAGCAGATTGTACTGGGTCTCAGTGCTCATACTATATTAGACCTTACTCTCAACTACGTTAACACTAATAGGTAACACCTATAGGTACAATTAAACAATGACTCTTCATTTCTTATCAATTAACAAAAGTCTTTTAATAACTATGTATATTTCAATACGAATTCATCAACAAGATCTAAAGGTTACCCTTTATTATCTTTTTTTATGGTTTGTAGCACGTGCAACATATAAGAGTCATATCGCTAGGATAAAAGAAGTCAAGTCGAATATTTCAATGAGAAGGCACATATTAGCATTATAACTAATGAGAGACCGAGCGGTATCAGCTCACTC |
| 10 | Pdhytk6 | TCAGAGCAGATTGTACTGGGTCTCAGTGCTTTTTGTGCTGTTGATAACAATTGATCGGAAAAAAGCCTCATGCTCATTTAAAGAATGTTCACTAAAGTTTGCCATAAAGTAGAAACTTCTTATAAAAACAGTCAGAAATATACAAAAAATGAGAGACCGAGCGGTATCAGCTCACTC |
| 11 | Pdhcho1 | TCAGAGCAGATTGTACTGGGTCTCAGTGCGATTTTCACGTGCGAATGATTGCTAAGAATTTTAATTTTGAGTATTTAAAGGATAAGCAATGACCAACGTATTTCATTTACTATTTTTTACTGATACGTGTAAGTGAATATTGGATATTAAGGGATCAATTGATTAAATTAGGTTTTATAAAAATATTGATTACAACTTGAAATATACTTTGTTAATATTTTCAAAATAAACTAGTTATTAGAACCAGAGAGCAATTTACAAACTCAAATTGCAAATTTAAGGTAGAATAATCTGAAAATATATTAAGTAACTGAGATTGATTTTGGAATAATGAGAGACCGAGCGGTATCAGCTCACTC |
| 12 | Pdhhta1 | TCAGAGCAGATTGTACTGGGTCTCAGTGCACCTATGTTTGTTTTGCTTATTTTGCGTAGTTATCACATACGCGTCGGTGATGTGTGATATGTAAACAAACTATAATTTACGCAGTAATTCTAAAATTTACGCGCGAATGGAGCAATACTGAAAATTTCCCATTTGCCTGTAAAAAATAGCATCTTCGATCAGCCCCCGTATCAGGTGGCTGTTAAATAAACAAACAGAATTTTGTTAAAGATATATAAATTTACACAATTCATCTTTCAATATAATAATTGGTAATTTTCCTTCTTGTTAATTAATCAATAATAATAATAATAAATACAAATGAGAGACCGAGCGGTATCAGCTCACTC |
| 13 | Pdhmrpl22 | TCAGAGCAGATTGTACTGGGTCTCAGTGCAATCAACGTGAATTCGTTGATGCGAGTAATAAGGAAGAGATTGTTGTATATCAGCGTTTATGAAATGCTTTGTTCGTCCCTTGATTTTTCCAACTATGAAAACTTTACAACCATTCGACTAAACAAATACTAACATCATCTAGTGGTTCAAATATCATGCACACACCAAACCTCATTTTGTTTGGGCCGTTGGTTTAGGATTATCAGCTCCATTGTTGTTGGCTATTTCTCCTATCAGAAGAAAGTACTTGTATTCTGACCATGAGCCAATTCCAAAAGTATACCCTTTACCAACCAGAGCCAGAGACACTAACTTAGCTGGTTTTGACGACGAGTAAGTATACAAACACGATAAATTATTGCCCGTTTAAGGGACGACAATAGAGCATTGGGCCATTACATTATTCCATTACATTACTATATTATTTAAGGTATTAATTGTTCATAAGAAGATGGTTAATATTTCATCGGAATGGCTCAAAAATTTCATTCAAATTTCTTACTTTACAAGACTCCGGTTACCACTCAACTAGTCATATCATAATATACGTTTATATTCTCTACATCACGCTGTAGCTTGCCTAGTATCCGCCATCTAGTAGTCATATAGGGCTTACATAAGGCCTAAAAACCGGTCGTGCCAAAACCAAAACAAATTTTGTCGCTTCGATTTAAATTTTCACCAAAAATGCTAACCAGACCATAATGGAAGGCCAGAACTAGGGTTAACTAAGGATAATGAGAGACCGAGCGGTATCAGCTCACTC |
| 14 | Pdhmyo4 | TCAGAGCAGATTGTACTGGGTCTCAGTGCAAAAGACCAAACACTCTTAAAACATATTAAACTAATTTATAAATTAACACTCTTCAGATCCATCAGGCGTATTCGATTACATACAACCACGTTCATATCTGATTAAAACTTGGATTTATTTAATATTTTAATAATCATTCTACCTTGAATCGTTACGAGTGTTTTTTTTTGAGTATTAACAAATTTTTTTTTTAATTTGTTGATTTATTCTGATTTATTCTGTTGAATTCATCTTATAGGTGCAGATTAGTTTCTGTTATATCAGCGGAGTATTTTTATTTCAAAGCATAATATATTGAGGCAACATTTGAAAATGAGAGACCGAGCGGTATCAGCTCACTC |
| 15 | Pdhpdc1 | TCAGAGCAGATTGTACTGGGTCTCAGTGCATAATAATTAGCTGAATGCGATTACGATCATGCGATTTCACGGAAAAACCGTGCGATTCGGATAATTAAGTTCACACGCAGCCAGTCATAAATTAGCTGCAAACTGGCTTGTAAAGTATATAAGAGGTTGGATCAGCATATAATTCAAGGAAATGTTAAACAGTATAATAAACATAATAATCATATTAATATAAAGAACAAATGAGAGACCGAGCGGTATCAGCTCACTC |
| 16 | Pdhmam33 | TCAGAGCAGATTGTACTGGGTCTCAGTGCGTATGCTTAATTGTTTTATAATTCTAATCATTTATCTAATATAAATCATATTTTGTTTGAAGCTAAAAATGTTAGGCAATTTTTTGTATCTTACAACCATATGATAGTTGTCGTAACAAACTTAACACTCAGTTACTCTGTGGTCCAGCGTATCCATCATAACGGGGTTTTTGCAGTACATAGTGCGTGTATTCACCAGATCTTTTTTAGCACATAGCTAGGCTTCTGTCCAGTTGAATATGGATAGCTAAAGATCACCAAGCTAATCAATTAGAAAAGGAATTAAATCAATCAACTAGTAATGAGAGACCGAGCGGTATCAGCTCACTC |
| 17 | Pdhrpl8 | TCAGAGCAGATTGTACTGGGTCTCAGTGCACACCTTAATTCTGGAATTTAGGTATAATATATATGACTAATCACAGTAATATTGTTGTGTAGTCTTTTACTGGCGACTTGCCTCTACAAATTCTCTTTATGGAGACATGAATTACGAAGCTGAATGTTTCTCTAACAATCACCCTCATTTGACGACTACAGTCTTTACTTGCTCGACTCCACCTTATTCTCCCATCTCCCGGTCAGTTTTTTTCCTTCTTTTTTGCTTATTACTTCACCTTTACCTTAACCCTCAGGTCATCACTATCGCCCCCATTACCAAAAAATAGACCTCCTCCGTAGTATACCCTTCTATTACTGCAAAATACGAAACGCAGTCACATCTCATCACATGATAAAATGGAATCGGTTTATGTACAGGAATCAAATCGTAGAATTTTTTTCTGTGACGAGCTTCTGCTCACATAGTGAAAAATTCAGCTATAATAAAAATTCACAATCTACAATCAGAGTTGCTAGTTCAAGTTATTATATATCAAAAATGAGAGACCGAGCGGTATCAGCTCACTC |
| 18 | Pdhrps3 | TCAGAGCAGATTGTACTGGGTCTCAGTGCGTTCATTGGTAGAGAGTCATTAACCATAAGTATAGTACTACAACTGACTTGTGCAGCATTTACTGGTAAAAAATTATTACATAACACGAACAAGACGATTACATAAACAAGATTGATTCAGAATTTTGGGGTCGTGGATCGTACCGAACATTTTTTTTCAGTTCACCAGGACGAAAAATTTTTCACATATAGGTGAAAATTATAGTTAAACTTCTTCTTGTTATATACAACAAATCATAAATATACAAAAAATGAGAGACCGAGCGGTATCAGCTCACTC |
| 19 | Pdhysa1 | TCAGAGCAGATTGTACTGGGTCTCAGTGCCGAATGTACACGAAGATCTTGCTTAAATAAGGAAATTATATATATGTGGGTATAGTCCTTTACACCTAGGTATTATATTCTTCTTTGAAATGTCCTGTAGAATCACTCAATATTCAAAATCGAGTATCAGAAATCTCACGAAACCGTCTTTTGCTGTGCATCAATATTATAAATACTTTTTCCACAACACTTCGAAAAAAAATGAGAGACCGAGCGGTATCAGCTCACTC |
| 20 | Pdhzuo1 | TCAGAGCAGATTGTACTGGGTCTCAGTGCGTGTTGTCAGATACTATTATCTAGACCAAATATAGCTCTTATAGATAGTTAGCATTCAACAAAATCCATATAGAGTGTAAACTAAATACCGGTACGCGTAATTTTAGGCACTCTATAAATCTCATTACATCCCAAAAATTTTTCAAGCATATCATGGCTAAAGGTGATAAGGTACTAATATTATTTAATATCGAATAAAAAATGAGAGACCGAGCGGTATCAGCTCACTC |
| 21 | Pdhcit2 | TCAGAGCAGATTGTACTGGGTCTCAGTGCCTGAACCAGTATGTAATTAGAGACTTTAGTGGAGCGGATGGGCCATAATATGAGCCATAATATGGGCCATAATGAATGCTTCCAGCGAATGATTCGACAAATACTAACTAGTACAGACTTTGAAAACTCGTTTATCTGAAATTTTACCAGAAAGAGCCGAAGAAGTTAAGCAATTCAAGAAGGAACACGGTAAAACCGTATGTATTATATTTGAACACAGTATGGTGTTGATGTAGTATCATTTAGACTATAGATGGACCAAAATGGCACCTGTTGATGGTTTTATCTGCCAACGGTACCTGAGGAATTTAATACCAGATTGTGAAAAATCGCGAATGCTCGATACTAACAAATATTTAGGTTATTGGTGAAGTCTTATTAGAACAAGCCTACGGTGGTAAATGAGAGACCGAGCGGTATCAGCTCACTC |
| 22 | Pdhcps1 | TCAGAGCAGATTGTACTGGGTCTCAGTGCTTGGTCTTGGGTATGTTTACGTACCAGTGACATTGACATTCCTATTGATCGTTATATGCGTCATACAACTATATTTTTAGCCTGGCATATTTGGTGCAATACATTAGTCTGTAGTACAAGTAAGAGTGCGGAATTATATCGGCTGGTGTATTTCCGAAGTCTCTCCATGTATAATGTCCAACTGCTATATTGACTGTAATATCACTTTTTGCTTTCAAATATAATACACTAATTCTCAGTATATACCTACTCTCCAACATTGTGCCGTGGTATGATCGGGTTTGAGAGTTGTCCTTGTTTTGGGCTTATCAGGTGAATAGATGGGGTCATTCGCTTATCAATTCCTAATTAGGTGCCCTACCCAATTTTTCAAAAATGGAATATAAGGGTGAACATCTACAGCGACTTCTTAGTACATTATTTGGAACAGAACCAATTGACAGAACCAATTGACAGAACCAATTGACAGAATCAATTGACAGAATCAATTGAAATATAAAAATGAGAGACCGAGCGGTATCAGCTCACTC |
| 23 | Pdhhxt2 | TCAGAGCAGATTGTACTGGGTCTCAGTGCTAATTTATTAATTTCTTGTATACCTATCTAGATCCATGCTTTAGTATATAAAATGATCTCGGAACATTAAGTAGATGCAGCCGATTAAAAATTATGTGTCCGCTATTTTTCTCCAATAAATCGTTGCCTCATAACTAAGCTGTCCTCAGATTTAGCTATGGTTTAGAACAATGTTCTATGTTGCATGGAGAAACTGGATGTTGATTCGCCTTTTCGCGAACTGAATAATCAGCCTTAGAATGGTATTGTCCCGCACCGGCACTTTTATGAGCGAGTGGGGCAACATGAATTTTTCCGGGGCATGGCACGGAGAAACTGGAGCGTGTATGGGATTAATCTTGCATGCCGGGATAAAAAATTTTTCCAGGTATTTTTCCGGAGGTTCTAAAATCTGTTATGCCTTGATGGTAGAAAGTATAAATAAGTGGGTTATCCCTCTTGAAAATTCAGAATTCTTTTCTTGTATTCAATTATTTATTTAATTTAGAAGTCAAATTATCAATGAGAGACCGAGCGGTATCAGCTCACTC |
| 24 | Pdhjen1 | TCAGAGCAGATTGTACTGGGTCTCAGTGCGCCAAAATGACAAGCAAATTCAACATGTACGTCACTTATCAAAAAGCTTATATTTTCTAAACTCCACAACAATTTGGGCGGCTCTGGATGAATTAGTTATCTGAATTACCGACAAGCATACATTATGGGTATATGGAGGTATTCTTTTTGGGGTTATCCCTTCTATATTGCACGAAGTGAGTTTTTTTTTTTTGGATCTAGTGCGATAATATGACAGGCTTTCAGGCCCCCAACCTACCCCACAAAAAATCTCTTTGCCGCAAAGTGAGTTACTACAAAAATACATAAGTAAGATATGTTATCCAGATATATGAGAATATTTGTAGAGTCGATAGTTATATCACTTAGAAAAATGAGAGACCGAGCGGTATCAGCTCACTC |

### Supplemental Table S5: *List of terminator sequences in the parts collection.*

| **S/N** | **Terminator** | **Nucleotide sequence** |
| --- | --- | --- |
| 1 | Tdhpfy1 | ATCAGAGCAGATTGTACTGGGTCTCATAAAAGGCCTGTCCGAATTCGGGCACTTGAATACCAATTTGCGTTATGTGGCCACGGGCTCTTTTCACGCTGTCATCACCATACTCTCGGTAACGGCTGTATGAGATGATGTATTATGTAGCGGGAAGTTTATAGTAACTAATATTATACAAGTATTGTAACTGTAGGGGTTTGCTACTCAAACTGTGACTACTATGTCATGATTATCGGCCTGATACCATATATATTACATTTTTACCACTCAATTCGCTCAATATATACATTATCTATAGACATTACATTCTGAAGGACTTATACCCTTGCAACGGTGCACCCTCTAGTGAAAAAACTGCAGAGCAAGCACTTCAAACAGGGTTTCTACCATTTTTGTAAGCCATTGAGAACCCATAGTCAATGAATCACTCCCTCAGAGACCGAGCGGTATCAGCTCACTC |
| 2 | Tdhfba1 | ATCAGAGCAGATTGTACTGGGTCTCATAAATTTTAACCCGTATAGAGCCATATATACTATATGTAATTTATCTATCTACCCTAATATAACGTCTCCAGTAAGAAAAGTTTTCCAGTGTGTGCGAATCAATTTTCTGATATCTAAAACCTCCTAAACGAATCAAATTATGTATGTAGCAATCCTCAGAGACCGAGCGGTATCAGCTCACTC |
| 3 | Tdhhhf2 | ATCAGAGCAGATTGTACTGGGTCTCATAAATTTTATACGATCAATATATGGGTTTATGTTATGAAAGGCATTGTAGGATGTAGAGTAGACTGGAGGTCTAAAATAGACTGGGGACGATATGAATAGGTTGATTAGGGGCCTGAAAATTGGAATGGTTTCTATGTGTTCTATATGGGTATTATATAGGTACTATTTGCTCTATTTGGGTATAATTTCGGCATGATATCTCTCCTCAGAGACCGAGCGGTATCAGCTCACTC |
| 4 | Tdhrpl28 | ATCAGAGCAGATTGTACTGGGTCTCATAAAATAAACTAAGTAATGAATTGTTTGTATATTTTACTCAAATAAATAATTGACTACGTGTTTCTTAATTCCGAGAGATTTTATTTCTTAATATAATTCACATTTTAGTGTTCAATGGATTGTAGGTACATGAATATCTATAGAGCTGACATTTAGTGAGCTTTTTGACGCAACAATACAAGAGAGGATATAGTTCCAAGCTTGTAGTGTTAGTCCTACTGTGACAAGACTAGAGTAAATTTAGAGGTATATAAAACCCAAATTACATACCATTTACTACAAGATAGTATCACCTACAAGCTGCCTCAGAGACCGAGCGGTATCAGCTCACTC |
| 5 | Tdhrps9 | ATCAGAGCAGATTGTACTGGGTCTCATAAAAATCATTTGATTTTTAGTTAATATTTACTTCACTTTAAATTAATAATATATTCTAATTTGGTCTTTTCCAATCTACTTTTCATTTATATTTATGTAGTATAGCAAAATTAATGGTTATTTATGGGTAATTTACTTTGTAATCTAGCGGATACTATCGTTACCATTCCCTATACTAAACGTACTCCCAATTGACCTATTGGGCACCATGCAACACTTATCACCATGTTCATGAGATCACTATAGCTATTACTCAATTAGGGTTCCAGCCCGATGTTACACAAAAACATCGAAAATGAAAATTCCTCAGAGACCGAGCGGTATCAGCTCACTC |
| 6 | Tdheno2 | ATCAGAGCAGATTGTACTGGGTCTCATAAAATCATTTAATGTCCTTATTTATATTTTGATAAAAGTCGTTTATTGAATACACGCTGCTAATAACTTGTTTATTTGTAGATCTATATTCCCTCAGAGACCGAGCGGTATCAGCTCACTC |
| 7 | Tdhhxt7 | ATCAGAGCAGATTGTACTGGGTCTCATAAATCCACTCTCGCGGACGAATACAATATAATAATATAATAATAATATAATATAATAGAACTCGGTAATAGAATAATTTGATCTCCCTTGAGGCGACCTATGTTGTGTATACGCTTTATGATATAGTTAATTAGATTGCATGTTATATACGCCTTATGATAAGTTCTTTGTAATATAGGGATGTATTTAATGCTGTTCAGGACTGTTAACCGCAGTAACTGATAATGCTATAGTGCTGAATACTAATGGGTTACAGTTAATGTAAAAGCCAAATTTTTGTTTTGTAGCCAGCAATCCGCTCATTAGCCAAATCAGGTAATCCGACCTTCGCTTTGTACTTAAAGAAGGCAATAACAATAAAGAACAAACGACTCAATCTCAATGAAATAAATTGCCCAAATACTACTTTTGTCTATCCAGCTTAACTACATAACATCTACATAAACTGTATTGCTGCAACTAGGAACATAACAACCATATCGGTGTATTATGTAATGTTTTTCCCTCAGAGACCGAGCGGTATCAGCTCACTC |
| 8 | Tdhhis5 | ATCAGAGCAGATTGTACTGGGTCTCATAAAGTGTATTATCTGTGGTATATATATATATATACATATAAACAAGTAAAGTTGTAGTATAGGTGATCCATAGCGTAATTAATTGTATGATAAAATTTTTGCCGGGTAACATTTTCGATTCTATGAGAAATTCAATTTTTTTCACCGACTTAACCTCAGAGACCGAGCGGTATCAGCTCACTC |
| 9 | Tdhytk6 | ATCAGAGCAGATTGTACTGGGTCTCATAAAATTGAACCGGGGGGATACATGGGTGTATTGATTCGAAGAATATCGATCGGTTTTTATTTAAGGTCGTCGTATTAGCTTTAGTAAATTTATTAATATACAAGTCTATTGAATAAATTAGTACTATAGCACCCATATATCAAATAGTGGGTGATTATTACTGTATTACATTTTTCAACAGTTACTATTTAATCGTAACGCATCCTCAGAGACCGAGCGGTATCAGCTCACTC |
| 11 | Tdheft2 | ATCAGAGCAGATTGTACTGGGTCTCATAAAGCTTATTAAATACTTAATTTTTAAATTATTAATGTTTGGTAAGTCAATTTCTATCTAATATGTATTTTTTTATTCTGTTTATTGAACTCTATTTGTAATTTGAATCTCTTTCAAGAATATATGATACTTTATAAAATCTAACACCCGTGTGACCACAACGCATAACGCCAATTGCTTTCGTTTATAGATTACCTCCTGTCCCTCAGAGACCGAGCGGTATCAGCTCACTC |
| 12 | Tdhhta1 | ATCAGAGCAGATTGTACTGGGTCTCATAAAGTGAATTAGATTTCATAATGAAGGGTTATATTTCTCTGTGTATTATATAAGTTTATTAATGTATATTATCTAGATGTGGGGTTTAATTTTCAATTTCATAGTCTTATTTAAAAAGAAGAACCTCAGAGACCGAGCGGTATCAGCTCACTC |
| 13 | Tdhmrpl22 | ATCAGAGCAGATTGTACTGGGTCTCATAAATCTTGTACATATATATATAATACTATAATTTATCACTATCATTTATTCCTGGACCTTATAAGACTATCGGTCGCTCGCTCACCGATGTATGATCCGATCACACCGTGTCTGTGCCTGCGATAACGTAATGTTTTTAGGTATTTCCGTCATGACCGAGTTTAACCCTATAAAAGACTCCTCTAACCGTCCACTGACCCTATTATTTCCGTGGCACAGTGACGTTATTCTAGCTCAATTTTCTTGTATATAGTATTAGTGCTCCAAAATGTCTAGACTACTTAATTCTTATCACCTGTTATACCTCAGAGACCGAGCGGTATCAGCTCACTC |
| 14 | Tdhpxr1 | ATCAGAGCAGATTGTACTGGGTCTCATAAATACAGATATAGATTACAATTTAATAAATTTATTACAACTATATATAAAATACATCCTCATTAATTCACTCGACCTCAGAGACCGAGCGGTATCAGCTCACTC |
| 15 | Tdhpdc1 | ATCAGAGCAGATTGTACTGGGTCTCATAAATAGAAAATGTTCGGTAATCTCAATTGAGACTGTATTTTTTAATTAATGATGCTTCTAATTAGAATAATACGAATATATATAGTCTCTATTTAATAAATCTTCTTATGGTAAGAAATAGAAATAATTGAATATAGTCAATTTTATTTCATTTAGAGTAAATTTCATAAATTTTGACATTGGTTGCTCGCATTTTGGTCCTTCCTCAGAGACCGAGCGGTATCAGCTCACTC |
| 16 | Tdhmam33 | ATCAGAGCAGATTGTACTGGGTCTCATAAATTAACTCCAAGTTCCTAAAGTTCTTTGACTAAGTCAAAATTACAATATCCAATTTTATGTGTTTTATATGCGTTTTACCTGTTAATAGTTCAAAGTTGTCTATAGTTTATGTTTAAAGTACGTTTTTAGCTGCAATATGTGTAAAAGTAAGGATGTCTATATCTCTATATATTGTGACTGCTTCTCGTGATCTGCGCATGCCTCAGAGACCGAGCGGTATCAGCTCACTC |
| 17 | Tdhrpl8 | ATCAGAGCAGATTGTACTGGGTCTCATAAAAATCTCATGTTTATAGTTATCTAATAATTATATATAATCGGTCGTTCCAAGTACTTTAATACATCTGAATCTACTTAATTTCTGGGGATAGTCTCTCGTATACTGTACCATATGTACTATATGTCAATGAAATTTTCAGCTGAGGTCCGGGTAATGAGGCTCGAATTTCTGGCCAATAACAGTTTTGATTTTTCTCATTTACCTCAGAGACCGAGCGGTATCAGCTCACTC |
| 18 | Tdhysa1 | ATCAGAGCAGATTGTACTGGGTCTCATAAAATGTGGTTATAAATATACTATAGATCACGATATATAGCTAGCTTTGAACGTTACTGATGATATTTGTTGTTTCTACGTAGTTAAATAAACCATCAAGACGCATTCTCTAAAATCCTCAGAGACCGAGCGGTATCAGCTCACTC |
| 19 | Tdhzuo1 | ATCAGAGCAGATTGTACTGGGTCTCATAAAGTCTATCATTAACTACGTTATTTATTAATTTTAGATACTGTTTGTATATAAATACATCTATACATTAACATGATATAATCGCTTCCAAAGACAACACCCCCATACTCCCTCAGAGACCGAGCGGTATCAGCTCACTC |
| 20 | Tdhcit2 | ATCAGAGCAGATTGTACTGGGTCTCATAAAGTTTGATGGTTTTTATCTCCTAACTTCCATGCCTATTTATTGTAATTTTTCCCATAAAGAAATGTTATTTATTAAAACTATTTGTTATTTATTTATGTATAGGTTCTTCTGTATGTAACGAATTGATGATTTTTATATATCTTTTTCCTCAGAGACCGAGCGGTATCAGCTCACTC |
| 21 | Tdhcps1 | ATCAGAGCAGATTGTACTGGGTCTCATAAATTAGAACGATAAATACATATTGGACATTTTTCACATCCAATCGTATCTTCTTCACATCCAATCATACCTTTTTGACTCCGCACTATCTCCTTTTGACTTGGCCTGCGAAGATCCATAGTTGAGCACTGACAATCTATTTCACCTGCGGAAGATGGGGCCGTCGATTCTCCGAGATAGCCAAAGCAATCAAATATGCCATACCTCAGAGACCGAGCGGTATCAGCTCACTC |
| 22 | Tdhhxt2 | ATCAGAGCAGATTGTACTGGGTCTCATAAATCCACTCTCGCGGACGAATACAATATAATAATATAATAATAATATAATATAATAGAACTCGGTAATAGAATAATTTGATCTCCCTTGAGGCGACCTATGTTGTGTATACGCTTTATGATATAGTTAATTAGATTGCATGTTATATACGCCTTATGATAAGTTCTTTGTAATATAGGGATGTATTTAATGCTGTTCAGGACTGTTAACCGCAGTAACTGATAATGCTATAGTGCTGAATACTAATGGGTTACAGTTAATGTAAAAGCCAAATTTTTGTTTTGTAGCCAGCAATCCGCTCATCCTCAGAGACCGAGCGGTATCAGCTCACTC |
| 23 | Tdhjen1 | ATCAGAGCAGATTGTACTGGGTCTCATAAATTCTGCATCACATACATTCATATAAGTTTAATATTAGTACTTCATGAAGCACTGGTTGTGATAACGGTACTAGATTGTGAGAACGGTGCGAATCTCATAATTGCAACATTTTTTGATTCTTCTATATATGTCCCAATTAGGAAGTAAGCGATATAGGTTTAACTTAATTCAAGTTTATTTATCAAATTATTGTAGAATGCCCTCAGAGACCGAGCGGTATCAGCTCACTC |
| 24 | Tdhmdh2 | ATCAGAGCAGATTGTACTGGGTCTCATAAAGATATGTAACTCCTTCCCTATATATGCACCACAATATTTATTTCGAACCGATTTTCTGACTTTTTTGTAGTTTTCCGTGTAAACTATTGATTTCCATGCAACTACTCCGGGTTTGAGTTGGAATATAGGTTTCGCCTCGTGGGTTTTCTTATCTCATCCTTTGTAGACAGGATTGGGTAATGTCATCAATATTCATACTTGAGATCCATTAGCTTCCTCCGCTGGCTCGGTGGGAGAACTAAATAACTACAACCTACTGCCTCTAGCAATGAGACTAAACAACATAGACCAATAGAGCAACCTCAGAGACCGAGCGGTATCAGCTCACTC |

### Supplemental Table S6: *DNA sequences of codon-optimized genes used in this study*

| GOI | Protein | Nucleotide sequence |
| --- | --- | --- |
| DHSAT | nourseothricin resistance | ATGAAAATTTCGGTGATCCCTGAGCAGGTGGCGGAAACATTGGATGCTGAGAACCATTTCATTGTTCGTGAAGTGTTCGATGTGCACCTATCCGACCAAGGCTTTGAACTATCTACCAGAAGTGTGAGCCCCTACCGGAAGGATTACATCTCGGATGATGACTCTGATGAGGACTCTGCTTGCTATGGCGCATTCATCGACCAAGAGCTTGTCGGGAAGATTGAACTCAACTCAACATGGAACGATCTAGCCTCTATCGAACACATTGTTGTGTCGCACACGCACCGAGGCAAAGGAGTCGCGCACAGTCTCATCGAATTTGCGAAAAAGTGGGCACTAAGCAGACAGCTCCTTGGCATACGATTAGAGACACAAACGAACAATGTACCTGCCTGCAATTTGTACGCAAAATGTGGCTTTACTCTCGGCGGCATTGACCTCTTCACGTATAAAACTAGACCTCAAGTCTCGAACGAAACAGCGATGTACTGGTACTGGTTCTCGGGAGCACAGGATGACGCCTAA |
| DhGFP | mVenus green fluorescent protein | ATGTCTAAAGGTGAAGAATTATTCACTGGTGTTGTCCCAATTTTGGTTGAATTAGATGGTGATGTTAATGGTCACAAATTTTCTGTCTCCGGTGAAGGTGAAGGTGATGCTACTTACGGTAAATTGACCTTAAAATTTATTTGTACTACTGGTAAATTGCCAGTTCCATGGCCAACCTTAGTCACTACTTTCGGTTATGGTGTTCAATGTTTTGCTAGATACCCAGATCATATGAAACAACATGACTTTTTCAAGTCTGCCATGCCAGAAGGTTATGTTCAAGAAAGAACTATTTTTTTCAAAGATGACGGTAACTACAAGACCAGAGCTGAAGTCAAGTTTGAAGGTGATACCTTAGTTAATAGAATCGAATTAAAAGGTATTGATTTTAAAGAAGATGGTAACATTTTAGGTCACAAATTGGAATACAACTATAACTCTCACAATGTTTACATCATGGCTGACAAACAAAAGAATGGTATCAAAGTTAACTTCAAAATTAGACACAACATTGAAGATGGTTCTGTTCAATTAGCTGACCATTATCAACAAAATACTCCAATTGGTGATGGTCCAGTCTTGTTACCAGACAACCATTACTTATCCACTCAATCTGCCTTATCCAAAGATCCAAACGAAAAGAGGGACCACATGGTCTTGTTAGAATTTGTTACTGCTGCTGGTATTACCCATGGTATGGATGAATTGTACAAATAA |
| MmTES | Mus musculus  Acyl-CoA Thioesterase | ATGTCAGCGCCTGAAGGTCTAGGTGATGCCCATGGGGATGCAGATCGTGGCGATCTTTCTGGCGACTTGCGTTCTGTTCTCGTAACTAGCGTATTAAACTTGGAACCATTAGACGAAGATCTTTACAGGGGTCGGCATTATTGGGTTCCTACATCTCAACGTTTATTTGGAGGTCAGATTATGGGTCAGGCTTTGGTTGCTGCTGCGAAATCTGTGTCTGAGGACGTACATGTCCATAGCTTACATTGTTATTTCGTACGTGCTGGTGATCCAAAAGTTCCCGTATTGTATCATGTTGAAAGAATAAGAACTGGAGCATCATTTTCTGTCCGTGCGGTTAAAGCCGTTCAACACGGAAAGGCTATTTTTATTTGCCAGGCTAGTTTCCAACAGATGCAGCCTAGTCCCCTGCAACATCAATTCTCTATGCCCAGTGTCCCACCGCCGGAAGATTTATTAGATCATGAAGCTTTAATCGATCAGTATTTGCGAGATCCTAATCTACATAAAAAATACAGAGTTGGATTGAACAGAGTAGCAGCTCAGGAAGTTCCTATCGAGATTAAGGTTGTTAATCCCCCAACTTTGACACAATTACAGGCTTTAGAACCTAAGCAAATGTTTTGGGTGAGAGCGCGTGGCTACATTGGTGAAGGTGATATCAAAATGCATTGCTGTGTAGCTGCCTACATAAGTGATTATGCTTTTCTTGGTACTGCCTTGTTACCACACCAATCGAAATATAAGGTAAATTTCATGGCTAGTTTAGATCATTCAATGTGGTTCCATGCTCCATTTAGAGCTGATCATTGGATGTTGTATGAATGCGAATCTCCATGGGCAGGTGGTTCTAGAGGCTTGGTCCATGGAAGATTGTGGCGGAGAGATGGAGTGTTAGCAGTAACCTGCGCTCAGGAGGGAGTGATTCGGCTCAAGCCTCAAGTCTCTGAATCTAAATTA |
| HpTES | Helicobacter pylori  Acyl-CoA Thioesterase | ATGAGATGCAGAGTGTATTACGAAGATACGGACTCGGAGGGGGTAGTGTACCATGCTAATTACTTGAAGTATTGCGAAAGAGCGAGATCTGAATTTTTCTTTAAACAAAATGTTTTGCCTGAGAATGAAGAGGGCGTGTTTGTTATTAGGTCGATTAAAGCTGATTTTTTTACACCAGCAAGTTTAGGCCAGGTTTTGGAAATCAGGACACAGATCAAGGAACTTAGGAAAGTTTTTGTAGTGTTGTTCCAGGAAATATACTGTATTCAAAATGCATCGCTTGAGCCGATGAAGCCATTTAAAGTTTTTGCATCGGAAATTAAATTTGGCTTCGTAAACAGGTCAACTTATTCCCCAATTGCCATCCCAAAGCTCTTTAAAGAATTATTAAATGCTATC |
| NiCAR | Nocardia iowensis  Carboxylic Acid Reductase | ATGGCGGTAGATTCACCGGATGAGAGACTACAACGTAGAATTGCTCAACTGTTTGCAGAAGATGAACAGGTTAAAGCAGCACGGCCATTGGAAGCAGTGTCTGCTGCAGTCTCAGCTCCTGGTATGAGATTGGCGCAGATTGCTGCAACAGTGATGGCAGGCTATGCAGACCGGCCAGCTGCCGGACAAAGAGCTTTCGAATTAAATACTGATGACGCAACTGGTCGGACTTCTCTAAGATTACTTCCTAGATTCGAGACGATCACATATCGTGAATTGTGGCAGAGGGTGGGTGAAGTGGCTGCCGCGTGGCACCATGATCCAGAAAACCCACTAAGAGCCGGGGATTTTGTGGCGCTTCTCGGGTTTACATCTATTGATTATGCGACATTAGACTTAGCTGACATTCACTTAGGAGCAGTTACTGTTCCTTTGCAGGCATCTGCTGCAGTAAGTCAATTGATTGCAATTTTGACCGAAACATCTCCTAGATTACTCGCGTCCACTCCAGAGCATCTCGACGCCGCTGTTGAATGTTTGTTGGCTGGAACAACACCTGAACGGTTAGTCGTTTTTGATTATCACCCAGAAGATGACGATCAAAGAGCAGCATTTGAATCGGCCCGTAGAAGATTAGCAGACGCGGGAAGTCTAGTAATAGTTGAAACTTTGGATGCTGTAAGGGCTAGAGGCCGTGATCTACCGGCAGCTCCCCTATTCGTTCCGGATACGGATGATGATCCGTTAGCATTGCTGATCTACACTAGTGGTTCTACAGGGACTCCGAAAGGTGCCATGTATACTAATCGTTTGGCCGCTACTATGTGGCAAGGAAATAGCATGCTCCAAGGTAATTCACAACGGGTAGGAATTAATCTTAACTACATGCCAATGTCACATATCGCAGGAAGAATTTCTCTTTTCGGAGTATTAGCTCGAGGTGGGACAGCCTACTTTGCGGCTAAGTCTGATATGTCGACATTGTTTGAGGACATTGGGCTAGTGAGGCCCACCGAAATCTTCTTTGTCCCAAGAGTATGCGACATGGTCTTTCAAAGATACCAGTCGGAGTTGGACCGTCGATCCGTAGCAGGTGCTGATTTGGATACTTTAGATAGAGAGGTGAAAGCTGACTTGAGACAGAATTACTTAGGTGGCAGGTTCTTAGTTGCTGTCGTAGGCAGTGCGCCATTGGCGGCAGAAATGAAAACTTTCATGGAATCGGTGCTAGATCTTCCATTGCACGACGGATATGGCTCAACAGAAGCTGGTGCCTCGGTCCTATTAGATAATCAGATCCAACGACCACCAGTTCTTGACTATAAGCTGGTAGATGTGCCTGAATTGGGTTATTTTAGAACCGATAGACCGCATCCACGAGGGGAATTATTACTTAAAGCCGAGACTACCATACCTGGTTACTATAAAAGACCAGAAGTAACCGCGGAAATTTTTGATGAAGATGGTTTTTACAAGACTGGAGACATAGTTGCGGAACTCGAACATGATCGTTTGGTGTATGTTGATAGAAGAAACAATGTTCTAAAGTTATCTCAGGGTGAATTTGTAACAGTTGCACATCTTGAAGCAGTGTTTGCATCTTCGCCATTGATTAGACAAATTTTCATCTATGGTTCCTCCGAAAGGTCGTATTTATTGGCCGTGATAGTTCCAACGGACGACGCTTTGAGAGGTAGAGATACTGCAACTTTAAAGTCCGCATTAGCTGAGTCTATCCAAAGAATTGCTAAGGATGCGAATTTGCAACCATACGAAATTCCACGTGATTTTCTTATTGAGACGGAGCCTTTCACAATAGCAAACGGATTGTTGTCCGGCATTGCCAAATTATTAAGACCAAACTTAAAGGAACGATATGGTGCTCAACTTGAGCAAATGTACACTGATCTTGCAACTGGTCAAGCTGATGAATTGCTGGCACTAAGAAGAGAAGCTGCTGACTTACCTGTATTGGAGACGGTGTCAAGAGCCGCTAAGGCTATGTTAGGTGTTGCCTCTGCCGATATGCGACCGGATGCACACTTTACTGATTTGGGTGGCGATTCTTTATCTGCTTTATCTTTCAGCAATCTGTTACATGAAATTTTTGGAGTGGAAGTTCCTGTAGGCGTCGTGGTATCGCCAGCGAATGAGTTAAGAGATCTAGCTAATTATATAGAAGCGGAGAGAAATTCCGGCGCAAAAAGGCCAACATTTACTTCTGTACATGGAGGGGGGTCTGAAATCAGAGCTGCTGACTTGACTCTAGATAAATTTATTGATGCAAGAACTTTGGCAGCCGCAGATTCAATTCCACATGCTCCCGTGCCTGCCCAAACCGTTTTATTAACAGGTGCCAATGGTTACTTAGGCAGATTCTTATGTTTGGAATGGTTAGAAAGATTAGATAAAACTGGAGGAACCTTAATATGTGTTGTACGAGGTTCAGACGCAGCAGCTGCCCGTAAACGATTAGATTCTGCATTTGACTCAGGTGATCCTGGACTATTAGAGCATTATCAACAATTAGCAGCTAGAACTTTGGAGGTGTTGGCTGGTGACATAGGGGATCCGAACTTAGGTTTGGATGATGCCACTTGGCAGAGGTTAGCGGAGACGGTTGACCTTATAGTACACCCGGCCGCTCTCGTAAATCATGTATTGCCTTATACACAACTTTTTGGTCCTAATGTAGTGGGAACTGCAGAAATCGTCAGACTCGCAATCACAGCAAGAAGGAAACCTGTTACTTATTTAAGTACTGTTGGAGTTGCTGATCAAGTGGACCCTGCTGAATATCAGGAGGATTCTGATGTTAGAGAAATGTCAGCGGTAAGAGTTGTTAGAGAATCTTATGCTAACGGATATGGTAATTCTAAGTGGGCAGGAGAAGTGTTACTTAGGGAAGCACACGATTTGTGCGGTTTACCAGTGGCAGTATTTAGATCTGATATGATTTTGGCTCATTCCCGTTATGCAGGTCAGTTAAATGTTCAGGACGTATTTACACGTCTGATCTTATCTTTGGTGGCCACAGGTATTGCTCCCTACTCATTCTATAGAACGGACGCTGATGGTAACAGACAAAGGGCACATTACGATGGTCTACCAGCTGATTTTACTGCAGCCGCCATAACTGCTTTGGGCATTCAGGCGACAGAAGGATTTAGAACTTATGACGTTCTGAACCCATACGATGATGGTATATCATTAGATGAATTCGTCGATTGGTTGGTTGAATCAGGACATCCAATTCAAAGAATAACTGATTATTCTGATTGGTTTCACCGTTTTGAGACAGCAATAAGAGCTTTGCCTGAAAAGCAACGGCAAGCTTCGGTTTTGCCCTTACTGGATGCTTACAGAAACCCGTGTCCGGCTGTTAGAGGTGCTATTTTGCCAGCTAAGGAATTTCAAGCTGCAGTCCAAACAGCAAAGATAGGTCCTGAACAAGATATACCACATTTGTCAGCACCATTAATCGATAAGTACGTATCAGACTTGGAGTTATTGCAATTGCTA |
| MmCAR | Mycobacterium marinum  Carboxylic Acid Reductase | ATGTCTCCAATCACTAGAGAAGAGAGATTGGAAAGAAGAATTCAAGACTTATACGCAAATGATCCTCAATTCGCAGCAGCAAAGCCGGCAACGGCAATTACTGCAGCTATTGAACGTCCAGGCTTACCATTACCTCAAATTATTGAAACCGTCATGACCGGATACGCTGATAGGCCAGCTTTGGCCCAAAGATCGGTTGAATTTGTCACAGATGCTGGGACAGGTCACACAACTTTACGACTATTACCACACTTCGAGACTATTTCTTATGGTGAGTTATGGGACCGTATATCGGCTCTTGCAGACGTCTTGTCTACAGAACAAACCGTCAAACCTGGTGATAGGGTTTGTTTACTGGGATTTAACAGTGTCGATTATGCAACAATTGATATGACATTGGCACGATTGGGGGCCGTAGCGGTACCGTTGCAAACTTCAGCAGCCATTACTCAATTACAGCCAATTGTAGCTGAAACTCAACCTACCATGATAGCAGCGTCTGTTGATGCATTAGCTGATGCCACTGAATTAGCCTTATCAGGTCAAACTGCAACAAGAGTCCTCGTGTTCGACCATCACAGACAAGTTGACGCTCATCGTGCAGCCGTGGAATCTGCGAGAGAAAGATTGGCGGGTTCTGCAGTTGTTGAAACTTTAGCCGAAGCAATAGCGAGAGGTGATGTTCCTAGGGGTGCTAGCGCTGGCAGTGCCCCAGGAACCGACGTGTCGGATGATTCTTTAGCCCTTCTAATTTATACCTCAGGATCTACAGGTGCGCCTAAAGGTGCAATGTACCCAAGGCGCAATGTGGCCACATTTTGGAGAAAGAGAACATGGTTCGAAGGTGGTTATGAACCATCAATTACTTTGAATTTTATGCCAATGTCTCATGTGATGGGTAGACAAATATTGTACGGGACCCTATGCAACGGCGGTACTGCATACTTTGTTGCAAAATCAGACTTGTCGACGCTTTTCGAGGATTTAGCATTGGTACGACCAACCGAACTCACTTTTGTTCCTAGAGTTTGGGACATGGTGTTCGATGAATTTCAATCTGAGGTGGATAGGAGATTGGTCGATGGAGCGGATAGAGTAGCTTTAGAAGCTCAAGTTAAAGCAGAGATTAGAAACGATGTTCTGGGCGGTAGATATACCTCTGCATTGACCGGTTCGGCACCTATTTCAGATGAAATGAAAGCTTGGGTCGAGGAGCTTTTGGATATGCATCTGGTCGAAGGATACGGTTCTACCGAAGCGGGGATGATCTTAATTGATGGTGCAATTCGTAGACCTGCAGTTTTGGATTACAAGCTGGTTGATGTTCCCGATCTGGGTTATTTTTTAACCGATAGACCACATCCTAGGGGTGAATTATTGGTTAAGACTGATTCCTTGTTTCCTGGTTATTACCAAAGAGCAGAAGTTACTGCTGATGTTTTTGATGCAGACGGTTTTTATAGGACCGGAGATATCATGGCCGAAGTCGGACCAGAGCAATTCGTCTATTTGGATCGTCGTAATAACGTCTTGAAATTGTCACAAGGAGAATTCGTTACCGTCAGTAAGTTGGAGGCCGTTTTTGGTGATTCGCCTTTGGTTAGACAAATATATATTTATGGCAACAGTGCGAGAGCATATTTGCTTGCAGTAATAGTACCAACTCAAGAAGCTTTGGACGCTGTACCGGTAGAGGAACTAAAAGCCAGATTGGGAGATTCATTACAAGAGGTTGCCAAAGCTGCTGGACTTCAATCCTATGAAATTCCCAGGGACTTCATAATCGAAACTACACCATGGACTTTGGAAAATGGATTACTAACCGGAATTAGGAAGCTCGCAAGACCACAATTAAAGAAACATTACGGTGAGCTTTTAGAGCAAATATACACAGATTTAGCACATGGACAAGCTGATGAGTTGCGCTCACTTAGACAGTCAGGAGCAGATGCACCTGTCTTGGTGACCGTTTGTAGAGCTGCAGCCGCCTTGTTAGGAGGCTCTGCATCAGATGTTCAACCAGACGCTCATTTTACAGACTTGGGGGGTGACTCATTAAGTGCTTTGAGCTTTACTAACTTATTACATGAGATTTTCGATATTGAAGTACCGGTTGGAGTCATCGTTAGCCCAGCAAATGATTTACAAGCTTTGGCTGATTATGTTGAAGCTGCGAGAAAACCAGGTTCTTCCCGTCCAACTTTCGCTAGTGTTCATGGGGCATCAAATGGTCAAGTTACAGAAGTCCATGCAGGTGATTTGTCGCTCGACAAGTTCATAGATGCCGCAACATTAGCTGAGGCCCCAAGGTTGCCCGCAGCTAATACCCAGGTACGTACCGTCTTGTTGACGGGAGCTACCGGTTTCTTAGGTAGGTACTTAGCCTTGGAATGGCTTGAAAGAATGGATCTCGTCGATGGTAAATTAATATGTTTAGTTAGAGCTAAATCGGACACCGAAGCCCGTGCCAGATTGGATAAGACTTTCGATAGTGGAGATCCAGAGTTGTTAGCTCATTACCGGGCTTTGGCTGGTGACCATCTTGAGGTACTAGCAGGCGACAAAGGTGAGGCAGATTTGGGTTTAGATAGGCAAACTTGGCAGAGATTGGCTGATACGGTTGATCTTATCGTTGATCCGGCCGCCTTGGTTAATCACGTATTACCCTACTCACAACTTTTCGGACCAAACGCATTGGGTACGGCTGAATTATTGCGGTTGGCGTTAACATCGAAGATCAAACCTTATTCCTATACTTCCACTATCGGTGTCGCTGACCAAATACCTCCTTCAGCTTTTACTGAGGACGCAGATATAAGAGTTATATCAGCCACTAGAGCAGTTGATGACTCGTATGCTAATGGATACAGCAACTCAAAATGGGCAGGCGAAGTACTTTTGCGTGAAGCCCATGACCTTTGTGGTTTACCAGTAGCTGTGTTTCGTTGTGATATGATTTTAGCCGATACAACATGGGCAGGTCAACTTAACGTCCCAGATATGTTTACTAGAATGATTTTGTCATTAGCTGCCACAGGTATTGCACCAGGTTCATTCTACGAATTAGCCGCCGATGGGGCTAGGCAAAGAGCTCATTACGATGGGTTGCCAGTTGAATTCATTGCTGAGGCCATTTCGACGCTCGGGGCCCAGTCCCAAGATGGTTTTCATACGTATCACGTTATGAACCCATACGACGATGGTATAGGTTTAGATGAATTCGTAGATTGGTTAAATGAGTCTGGTTGCCCAATTCAAAGAATTGCAGATTATGGAGATTGGTTGCAAAGATTTGAGACGGCACTCAGAGCCCTACCTGATAGACAACGACACTCTTCTCTATTACCACTCCTTCATAATTACCGGCAACCTGAACGGCCAGTCAGAGGCTCGATTGCTCCAACTGATAGATTCAGAGCTGCTGTTCAGGAAGCTAAAATTGGTCCAGATAAGGATATTCCTCATGTAGGAGCACCAATTATTGTAAAGTATGTTTCGGATTTAAGACTGTTAGGATTACTG |
| CmCAR | Clavibacter michiganensis  Carboxylic Acid Reductase | ATGGGTAGTGAAAATATGGGAACCGCACAGATGAGATCTCAGCATGATGACACGAGTATAGAAGCCATTTTTGAACAACATGCCCAAAGAACTGCGTTAAGACAACGATCTGGTCCAGAGATCACTGATACTTCTTTCAGAGAGCTCTGGGATCGTGCTGCAGCACTCGCTGCGGCATTAGGTGAAACTGTTTCGGCAGGTGACAGAATTGCCGTTTTGGGTACCGCAACTGCTGACGCAGTTACTTTGGATTTAGCTACTTGGATATTGGGTGCAGTTTCTGTGCCATTACAGGCTAGTGCTCCTGTTGGTGCCCTTTCAGCCATTGTTGAAGAAACCACTCCCGTCTGGATAGCTGCTACCGCAGAACAATCAGCTACTGCACGAGCTGTTGTAGAAGCATCTGCTGACGGAATCAGAACCATGTTACTCGACACAGGTACAGGAACAGGAACTGATACTGATCTTACTTTGGAAGCGTTAGTTGCGAGGGGAGCAGGGTTACCGAGAAGATCGCCATGGCATCCCGCACCAGGAGATGACCCGTTAGCTTTATTATTGTACACGTCGGGAAGCACCGGAACTCCTAAAGGTGCTATGTATACCAGGTCTATGGTAGAAAGAATGTGGCATGCATTAAGACCAGACCCCGCGGCAGGGGCAGACGCCGCAGACGCTGCTGACGCTATTGTAGGTTATGCGTACTTGCCTATGTCGCATTTGACGGGAAGATCCTCACTATTGGCGACATTAGGTCGTGGTGGAACTGTAGCTTTAGCAACCTCGACCGATTTGTCCACGTTATTCGATGACTTAAGAGCATTCGCCCCAACTGAATTCGTATTCGTCCCTAGAGTAGCTGAAATGGTCAGGCAAGAAGGTGATAGAGAGGAACAAAGAAGATTGGCTGCTGGTGGCACAGACCCTGATGCGGTAAGAGCAGCTGTGCAGGCCGACTTAAGAGTAAGAGCATTCGGTGGGAGAATTAGTCGTGCGATATGTACTTCGGCCCCACTTACACCAGAACTTCGTACTTATATAGAGGGATGTTTAAGAATAACTCTTCATGATTTATACGGCTCAACTGAAGCTGGTGGTATTTTACATGATGGGGTAATTCAACAACCCCCAGTTACTGAACATAAGCTGGTAGATGTTCCCGAACTCGGCTATAGGACTACTGATTTACCACACCCCAGGGGAGAATTATTAATCAAGTCGACCGCAGTTATTGCCGGTTACTTCAGACGCCCAGATGTTACAGCTGCAGTATTTGATGAGGATGGTTTCTATCGTACGGGGGATGTTATGGCGCAAACGGGACCAGATACCTACGAATACCTTGATAGGAGAAATAATGTTCTAAAATTGTCTCAAGGTGAATTCGTTGCTGTTGCATCCTTGGAAGCCACCTATGGGGGTACACCAGAAGTTCATCAAATCGCATTGCACGGTGATTCAAGACACGCATTCTTAGTGGCAGTTGTAGTCCCTGCTGATGCAGGGGCCTCAGACAGAGATGTTTTGGCCGCGTTACAGAGAACCGCCAGAGAACAAGGCCTTGCCCCTTATGAGGTGCCAAGGGGTGTTATTGTGGAACCTAGGCCTTTCACAGTAGACGATGGTATGTTGTCAGACGCTGGGAAGCTCTTAAGGCTCAGACTAACTCAAAGATATGGGGAGAGATTTGCAGCCTTGTATGATGCGTTGGAGGAACAACAAACTGGATCGCTCGTTGCAGCTTTAAGAGACAGAGCTGGAGACGAACCAACAGTTGATACTGTAGTTAGAGCTGCATTACAACTTTTAGGTGCCGAAGTGTCGCCAGCAACTGCTGCCGCGGCAAGATTTTCTGACTTGGGTGGAGACTCTTTGAGTGCATTAACCTTCTCAGGTATTTTAGAGGATGTGTTCGGTACTGAAGTGCCTGTTGGTGTTTTAACAGATCCAACAAATGATTTGGCTGCTGTGGCTGCATATGTCGATCGAAGTGCGTCCGACGATCGACCAACAGTCACTAGAGTTCATGGTGCTTCAGCTTCGACGTTACGTGTGGATGATTTGAGACTTGACCGTGTTCTGGGTGCTATTCCAACCCCAGTCCCTCGTGCATCAGCTAGTAGACCAGGGTCTAGGACTGTCTTACTCACTGGTGCAAATGGATACCTTGGTAGATTTATGGCAATTGATTGGTTGGAGAGGTTGGCTCCAGAAGGTGGGACGTTAGTTTGTGTGGTAAGAGGCGCTGATGACGCCGATGCTAGGAGACGATTAGAAGCAGCCTTTGCAGCTGATCCAGCGTTCGCAAGGCGCTTCGCCGAATTGGCTGGTAGTTTAGAGGTAATTGCAGGTGATGTGTCCGAACATAGATTGGGCTTAGATGATGCAAGATGGACAGACTTGGCTGCTAGAGTAGATTTAGTAGCTCATGCAGCTGCTCTTGTTAATCATGTGTTGCCATACTCTGCACTTTTCGGACCAAACGTTGTTGGAACTGCCGAAGTTATCAGGCTTGCCATTGCTGCAGGTTCAGTACCGGTTACTTTTGTTAGCTCGGTGGCCGTTGCTGGAGGTGCAAGACCATCAGCTGCAGCGGATGCTGAACCTAGTGCTCCAGGAGCTTTAGATGAAGATGCTGATATCAGAGCAACTATACCAGAGTGGGCCGTAGGAGATGAATATGCCAATGGTTATGGAGCATCTAAATGGGCATCTGAAGTCCTTTTAAGAGAAGCTTATGAACACCATGGAGTACCTGTAGCTGTTTTTAGATCGGATATGATCCTTGCACATCCAAGATGGAGAGGTCAAGTTAATCTTCCAGATGTTTTTACTCGGCTTATATGGAGTGTTTTAACGACAGGGCTTGCTCCCGCTTCCTTCGTTAGGCGTGGTCCTGATGGAGAAAGACAAAGATCTCATTATGACGGACTACCAGCTGACTTTACGGCCGCAGCGATTGATGGTATCGGGGCTGCCATCACTGAAGGTCATAGGACTTTCAATGTCGTGAACCCTCATGATGATGGTGTCTCATTAGATACATTTGTTGATTGGCTAAGAGAGGATGGTCATGACATAGAAAGAGTCGAAGATCATGCTGAATGGGTTGAGAGATTTAGAGATGCCTTAGCTTCATTGCCTGACGCAGACCGGGCTAGAAGCGTTTTGCCCTTAATGCATGCTTTTGCTGCACCAGAAGAAACTCACGCCGGATCAGCAATACCCGCCGATGCATTTGCTGCTGCTGTTAGGGAGGTTAGGCCATTAGGGGCATCGGAAATTCCGAGTTTGGATCGTGCTTTAATAGCAAAAGTTGCTGATGATTTGGCTTTTCTTGGCTTACTCGAACCTGCTAGGGCGGCGGCAGCC |
| PmADO | Prochlorococcus marinus  Aldehyde Decarbonylase | ATGCCAACCTTGGAAATGCCAGTCGCTGCTGTTTTAGATTCCACAGTTGGCTCGAGTGAAGCGCTTCCAGATTTCACGTCTGATAGGTATAAAGACGCATATTCTAGAATAAATGCAATTGTTATAGAGGGCGAGCAAGAAGCCCATGATAATTACATAGCGATTGGGACGTTGCTTCCCGACCACGTGGAAGAATTAAAGAGATTAGCTAAAATGGAAATGAGGCATAAAAAAGGATTCACCGCCTGTGGAAAAAACTTAGGGGTTGAAGCTGACATGGATTTTGCCAGGGAGTTCTTTGCACCGCTCAGAGACAACTTTCAAACCGCTTTGGGTCAAGGCAAGACTCCTACTTGTTTATTAATACAAGCGTTACTGATTGAGGCTTTTGCCATTTCAGCTTACCATACCTATATTCCAGTTAGTGATCCTTTTGCCAGAAAAATAACCGAAGGTGTTGTCAAAGACGAGTATACACACTTGAATTATGGAGAAGCATGGTTAAAAGCTAACTTGGAAAGTTGTCGTGAGGAATTATTGGAAGCGAATAGAGAAAATCTTCCATTGATCAGAAGAATGTTAGATCAAGTGGCCGGGGACGCTGCTGTTTTACAAATGGATAAAGAGGACTTGATTGAAGATTTTCTTATTGCTTATCAAGAATCACTTACCGAAATTGGATTTAATACTAGAGAGATTACAAGGATGGCTGCAGCTGCATTGGTATCA |
| NpADO | Nostoc punctiforme  Aldehyde Decarbonylase | ATGCAGCAATTAACTGATCAATCAAAAGAATTGGATTTCAAATCAGAAACTTATAAAGATGCATACTCCAGAATTAATGCCATCGTTATTGAAGGCGAACAAGAGGCACACGAGAATTATATCACATTGGCTCAGTTATTGCCAGAAAGTCATGACGAGCTAATTAGGTTGTCGAAAATGGAGTCTAGGCATAAGAAAGGATTTGAAGCATGTGGTAGGAACCTAGCTGTAACACCTGATTTGCAATTTGCAAAGGAATTTTTTTCCGGTTTGCATCAAAATTTCCAAACAGCAGCAGCGGAGGGGAAAGTGGTTACATGTTTATTAATCCAAAGTTTAATTATTGAATGTTTTGCTATTGCCGCTTATAATATTTACATTCCTGTAGCTGATGATTTTGCGCGTAAAATAACTGAGGGGGTCGTAAAAGAAGAATACTCACATTTAAATTTCGGCGAAGTATGGTTGAAGGAGCATTTTGCTGAATCTAAGGCTGAGTTAGAATTAGCTAACCGTCAAAATTTACCAATTGTGTGGAAGATGCTCAACCAAGTTGAAGGTGACGCTCACACAATGGCAATGGAAAAAGACGCATTGGTTGAGGATTTTATGATACAATACGGAGAGGCGTTATCTAATATTGGATTCTCAACAAGAGATATAATGCGTTTGTCTGCATATGGTTTGATTGGTGCG |
| AtADO | Atelocyanobacterium thalassa   Aldehyde Decarbonylase | ATGCAGGAGTTAGCTTTACGCTCTGAACTTGACTTCAATTCAGAAACTTACAAAGATGCTTACTCTCGTATAAATGCAATTGTGATAGAGGGTGAACAAGAAGCATATCAAAACTATTTAGATATGGTTCATATGTTACCAAAAAATAAGGACGAACTAGTACGATTATCGAAAATGGAAAATAGACACAAGACTGGGTTCCAGGCATGTGGGAAGAATTTAAATGTAATTCCAGATATGCAGTACGCTAAGGAGTTCTTTTCGCAATTACACGAGAATTTCCAGATCGCTAAAAATGAAAAAAAAGTTGTTACATGCTTATTGATTCAGGCTTTGATAATAGAAGCATTCGCAATTGCCGCTTACAACATTTACATTCCCGTAGCAGATCCTTTTGCCAGAAAAATAACTGAAAATGTAGTAAAAGACGAATATAAACACCTTAATTTTGGCGAGGTTTGGCTTGGAGAGAATTTTGAATCGTCAAAAATTGAATTAGAAGAGGCAAATAAGACCAATTTACCTATAGTATGGAAGATGTTAAATGAAGTCGAACAAGATGCATCAATTTTAGGCATGGAAAAGGAAGCCCTAGTTGAAGATTTCATGATAAGTTATGGTGAGGCTCTAGGTAATATAGGTTTCTCAACTCGAGAAATAATGCGAATGTCCAGCCATGGTCTTAGAGCCTCG |
